# Stochastic processes dominate the within and between host evolution of influenza virus

**DOI:** 10.1101/176362

**Authors:** John T. McCrone, Robert J. Woods, Emily T. Martin, Ryan E. Malosh, Arnold S. Monto, Adam S. Lauring

## Abstract

The global evolutionary dynamics of influenza virus ultimately derive from processes that take place within and between infected individuals. Here we define the dynamics of influenza A virus populations in human hosts through next generation sequencing of 249 specimens from 200 individuals collected over 6290 person-seasons of observation. Because these viruses were collected over 5 seasons from individuals in a prospective community-based cohort, they are broadly representative of natural human infections with seasonal viruses. We used viral sequence data from 35 serially sampled individuals to estimate a within host effective population size of 30-70 and an in vivo mutation rate of 4x10^−5^ per nucleotide per cellular infectious cycle. These estimates are consistent across several models and robust to the models' underlying assumptions. We also identified 43 epidemiologically linked and genetically validated transmission pairs. Maximum likelihood optimization of multiple transmission models estimates an effective transmission bottleneck of 1-2 distinct genomes. Our data suggest that positive selection of novel viral variants is inefficient at the level of the individual host and that genetic drift and other stochastic processes dominate the within and between host evolution of influenza A viruses.

## Introduction

The rapid evolution of influenza viruses has led to reduced vaccine efficacy, widespread drug resistance, and the continuing emergence of novel strains. Broadly speaking, evolution is the product of deterministic processes, such as selection, and stochastic processes, such as genetic drift (Kouyos et al. 2006). The relative contribution of each is greatly affected by the effective population size, or size of an idealized population whose dynamics are similar to that of the population in question (Rouzine et al. 2001). If the effective population size of a virus is large, as in quasispecies models, evolution is largely deterministic and the frequency of a mutation can be predicted based on its starting frequency and selection coefficient. In small populations, selection is inefficient, and changes in mutation frequency are strongly influenced by migration or genetic drift.

Viral dynamics may differ across spatial and temporal scales, and a complete understanding of influenza evolution requires studies at all levels (Nelson & Holmes 2007; Holmes 2009). The global evolution of influenza A virus (IAV) is dominated by the positive selection of novel antigenic variants that circulate in the tropics and subsequently seed annual epidemics in the Northern and Southern hemisphere (Rambaut et al. 2008). Whole genome sequencing has also demonstrated the importance of intrasubtype reassortment to the emergence of diverse strains that differ in their antigenicity. While continual positive selection of antigenically drifted variants drives global patterns, whole genome sequencing of viruses on more local scales suggests the importance of stochastic processes such as strain migration and within-clade reassortment (Nelson et al. 2006).

It is now feasible to efficiently sequence patient-derived isolates at sufficient depth of coverage to define the diversity and dynamics of virus evolution within individual hosts (Kao et al. 2014). Studies of IAV populations in animal and human systems suggest that most intrahost single nucleotide variants (iSNV) are rare and that intrahost populations are subject to strong purifying selection (Rogers et al. 2015; Murcia et al. 2010; Iqbal et al. 2009; Poon et al. 2016; Dinis et al. 2016; Debbink et al. 2017). While positive selection of adaptive variants is commonly observed in cell culture (Doud et al. 2017; ARCHETTI & HORSFALL 1950; Foll et al. 2014), it has only been documented within human hosts in the extreme cases of drug resistance (Gubareva et al. 2001; Ghedin et al. 2011; Rogers et al. 2015), long-term infection of immunocompromised hosts (Xue et al. 2017) or experimental infections with attenuated viruses (Sobel Leonard et al. 2016). Indeed, we and others have been unable to identify evidence for positive selection in natural human infections (Debbink et al. 2017; Dinis et al. 2016), and its relevance to within host processes is unclear.

Despite limited evidence for positive selection, it is clear that novel mutations do arise within hosts. Their potential for subsequent spread through host populations is determined by the size of the transmission bottleneck (Alizon et al. 2011; Zwart & Elena 2015). If the transmission bottleneck is sufficiently wide, low frequency variants can plausibly be transmitted and spread through host populations (Geoghegan et al. 2016). Because the transmission bottleneck is conceptually similar to the effective population size between hosts, its size will also inform the relative importance of selection and genetic drift in determining which variants are transmitted. While experimental infections of ferrets suggest a very narrow transmission bottleneck (Varble et al. 2014; Wilker et al. 2013), studies of equine influenza support a bottleneck wide enough to allow transmission of rare iSNV (Hughes et al. 2012; Murcia et al. 2010). The only available genetic study of influenza virus transmission in humans estimated a remarkably large transmission bottleneck, allowing for transmission of 100-200 genomes (Poon et al. 2016; Sobel Leonard et al. 2017).

Here, we use next generation sequencing of within host influenza virus populations to define the evolutionary dynamics of influenza A viruses (IAV) within and between human hosts. We apply a benchmarked analysis pipeline to identify iSNV and to characterize the genetic diversity of H3N2 and H1N1 populations collected over five post-pandemic seasons from individuals enrolled in a prospective household study of influenza. We use these data to estimate the *in vivo* mutation rate and the within and between host effective population size. We find that intrahost populations are characterized by purifying selection, a small effective population size, and limited positive selection. Contrary to what has been previously reported for human influenza transmission (Poon et al. 2016), but consistent with what has been observed in other viruses (Zwart & Elena 2015), we identify a very tight effective transmission bottleneck that limits the transmission of rare variants.

## Results

We used next generation sequencing to characterize influenza virus populations collected from individuals enrolled in the Household Influenza Vaccine Effectiveness (HIVE) study (Monto et al. 2014; Ohmit et al. 2013; Ohmit et al. 2015; Ohmit et al. 2016; Petrie et al. 2013), a community-based cohort that enrolls 213-340 households of 3 or more individuals in Southeastern Michigan each year (Table 1). These households are followed prospectively from October to April, with symptom-triggered collection of nasal and throat swab specimens for identification of respiratory viruses by RT-PCR (see Methods). In contrast to case-ascertained studies, which identify households based on an index case who seeks medical care, the HIVE study identifies symptomatic individuals regardless of illness severity. In the first four seasons of the study (2010-2011 through 2013-2014), respiratory specimens were collected 0-7 days after illness onset. Beginning in the 2014-2015 season, each individual provided two samples, a self-collected specimen at the time of symptom onset and a clinic-collected specimen obtained 0-7 days later. Each year, 59-69% of individuals had self-reported or confirmed receipt of that season's vaccine prior to local circulation of influenza virus.

**Table 1.** Influenza viruses over five seasons in a household cohort.

|  | 2010-2011 | 2011-2012 | 2012-2013 | 2013-2014 | 2014-2015 |
| --- | --- | --- | --- | --- | --- |
| Households | 328 | 213 | 321 | 232 | 340 |
| Participants | 1441 | 943 | 1426 | 1049 | 1431 |
| Vaccinated, n (%)<sup>a</sup> | 934 (65) | 554 (59) | 942 (66) | 722 (69) | 992 (69) |
| IAV Positive Individuals<sup>b</sup> | 86 | 23 | 69 | 48 | 166 |
| H1N1 | 26 | 1 | 3 | 47 | 0 |
| H3N2 | 58 | 22 | 66 | 1 | 166 |
| IAV Positive Households<sup>c</sup> |  |  |  |  |  |
| Two individuals | 13 | 2 | 9 | 7 | 23 |
| Three individuals | 5 | 2 | 3 | 3 | 11 |
| Four individuals | - | - | 1 | 2 | 4 |
| High Quality NGS Pairs<sup>d</sup> | 4 | 1 | 2 | 6 | 39 |

<sup>a</sup> Self reported or confirmed receipt of vaccine prior to the specified season.
<sup>b</sup> RT-PCR confirmed infection.
<sup>c</sup> Households in which two individuals were positive within 7 days of each other. In cases of trios and quartets, the putative chains could have no pair with onset >7 days apart.
<sup>d</sup> Samples with >10<sup>3</sup> genome copies per µl of transport medium, adequate amplification of all 8 genomic segments, and average sequencing coverage >10<sup>3</sup> per nucleotide.

Over five seasons and nearly 6,290 person-seasons of observation, we identified 77 cases of influenza A/H1N1pdm09 infection and 313 cases of influenza A/H3N2 infection (Table 1). Approximately half of the cases (n=166) were identified in the 2014-2015 season, in which there was an antigenic mismatch between the vaccine and circulating strains (Flannery et al. 2016). All other seasons were antigenically matched. Individuals within a household were considered an epidemiologically linked transmission pair if they were both positive for the same subtype of influenza virus within 7 days of each other. Several households had 3 or 4 symptomatic cases within this one-week window, suggestive of longer chains of transmission (Table 1).

### Within host populations have low genetic diversity

We processed all specimens for viral load quantification and next generation sequencing. Viral load measurements (genome copies per μl) were used for quality control in variant calling, which we have shown is highly sensitive to input titer (McCrone & Lauring 2016) (Figure 1A). Accordingly, we report data on 249 high quality specimens from 200 individuals, which had a viral load of >10^3^ copies per microliter of transport media, adequate RT-PCR amplification of all eight genomic segments, and an average read coverage of >10^3^ across the genome (Table 1, Supplementary Figure 1).

**Figure 1.**
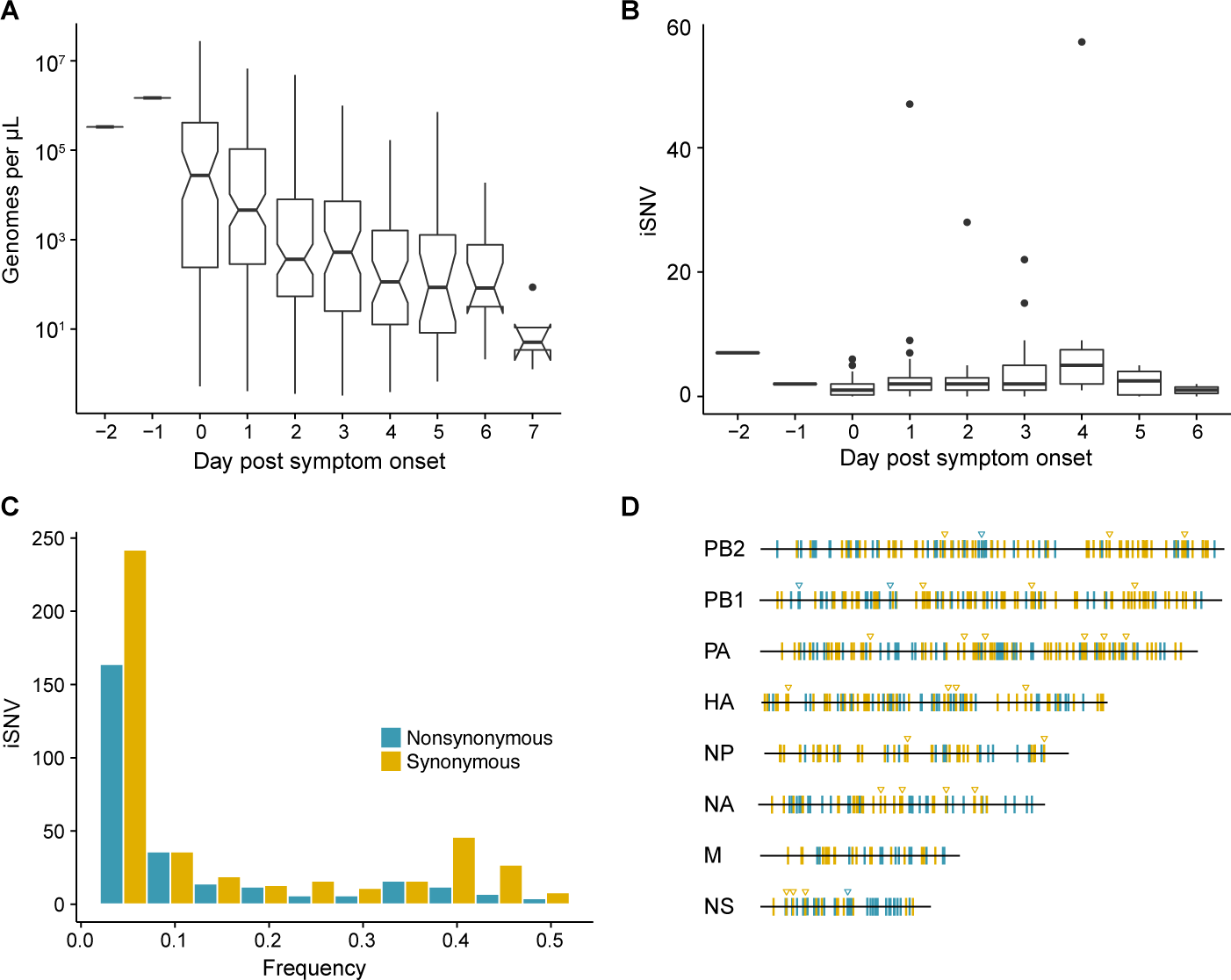
Within host diversity of IAV populations. (A) Boxplots (median, 25th and 75th percentiles, whiskers extend to most extreme point within median ± 1.5 x IQR) of the number of viral genomes per microliter transport media stratified by day post symptom onset. Notches represent the approximate 95% confidence interval of the median. (B) Boxplots (median, 25th and 75th percentiles, whiskers extend to most extreme point within median ± 1.5 x IQR) of the number of iSNV in 249 high quality samples stratified by day post symptom onset. (C) Histogram of within host iSNV frequency in 249 high quality samples. Bin width is 0.05 beginning at 0.02. Mutations are colored nonsynonymous (blue) and synonymous (gold) (D) Location of all identified iSNV in the influenza A genome. Mutations are colored nonsynonymous (blue) and synonymous (gold) relative to that sample's consensus sequence. Triangles signify mutations that were found in more than one individual in a given season.

We identified intrahost single nucleotide variants (iSNV) using our empirically validated analysis pipeline (McCrone & Lauring 2016). Our approach relies heavily on the variant caller DeepSNV, which uses a clonal plasmid control to distinguish between true iSNV and errors introduced during sample preparation and/or sequencing (Gerstung et al. 2012). Given the diversity of influenza viruses that circulate locally each season, there were a number of instances in which our patient-derived samples had mutations that were essentially fixed (>0.95 frequency) relative to the clonal control. DeepSNV is unable to estimate an error rate for the control or reference base at these positions. We therefore performed an additional benchmarking experiment to identify a threshold for majority iSNV at which we could correctly infer whether or not the corresponding minor allele was also present (see Methods). We found that we could correctly identify a minor allele at a frequency of ≥2% when the frequency of the major allele was ≤98% We therefore report data on iSNV present at frequencies between 2 and 98%. As expected, this threshold improved the specificity of our iSNV identification and decreased our sensitivity to detect variants below 5% compared to our initial validation experiment (McCrone & Lauring 2016), which did not employ a frequency threshold (Supplementary Table 1).

Consistent with our previous studies and those of others, we found that the within host diversity of human influenza A virus (IAV) populations is low (Poon et al. 2016; Dinis et al. 2016; Debbink et al. 2017; Sobel Leonard et al. 2016; McCrone & Lauring 2016). Two hundred forty-three out of the 249 samples had fewer than 10 minority iSNV (median 2, IQR 1-3). There were 6 samples with greater than 10 minority iSNV. In 3 of these cases, the frequency of iSNVs were tightly distributed about a mean suggesting that the iSNV were linked and that the samples represented mixed infections. Consistent with this hypothesis, putative genomic haplotypes based on these minority iSNV clustered with distinct isolates on phylogenetic trees (Supplementary Figures 2 and 3). While viral shedding was well correlated with days post symptom onset (Figure 1A) the number of minority iSNV identified was not affected by the day of infection, viral load, subtype, or vaccination status (Figure 1B and Supplementary Figure 4).

The vast majority of minority variants were rare (frequency 0.02-0.07), and iSNV were distributed evenly across the genome (Figure 1C and 1D). The ratio of nonsynonymous to synonymous variants was 0.64 and was never greater than 1 in any 5% bin, which suggests that within host populations were under purifying selection. We also found that minority variants were rarely shared among multiple individuals. Ninety-five percent of minority iSNV were only found once, 4.7% were found in 2 individuals, and no minority iSNV were found in more than 3 individuals. The low level of shared diversity suggests that within host populations were exploring distinct regions of sequence space with little evidence for parallel evolution. Of the 31 minority iSNV that were found in multiple individuals (triangles in Figure 1D), 4 were nonsynonymous.

Although the full range of the H3 antigenic sites have not been functionally defined, it is estimated that 131 of the 329 amino acids in HA1 lie in or near these sites (Lee & Chen 2004). We identified 17 minority nonsynonymous iSNV in these regions (Supplementary Table 2). Six of these were in positions that differ among antigenically drifted viruses (Smith et al. 2004; Wiley et al. 1981), and two (193S and 189N) lie in the “antigenic ridge” that is a major contributor to drift (Koel et al. 2013). Three of these have been detected at the global level as consensus variants since the time of isolation (128A, 193S and 262N) with two (193S and 262N) seemingly increasing in global frequency (Neher & Bedford 2015) (Supplementary Figure 5). Additionally, we identified 1 putative H1N1 antigenic variant (208K in C_a_) (Caton et al. 1982; Xu et al. 2010). In total, putative antigenic variants account for 1.0-2.5% of minority iSNV identified and were found in 3.5-8.0% of infections. None of these iSNV were shared among multiple individuals.

### Estimation of effective population size

Given the above observations, we hypothesized that within host populations of IAV are under purifying selection and that variants that rise to detectable levels do so by a neutral process as opposed to positive selection. Consistent with this hypothesis, we found that nonsynonymous and synonymous iSNV exhibited similar changes in frequency over time in the 35 individuals who provided serial specimens that contained iSNV (Figure 2A and 2B). We used a maximum likelihood approach to estimate the within host effective population size (N_e_) of IAV by fitting a diffusion approximation of the Wright-Fisher model (Kimura 1955). This model assumes that changes in iSNV frequency are due solely to random genetic drift and not selection, that iSNV are independent of one another, and that the effective population is sufficiently large to justify a continuous approximation to changes in allele frequency. The diffusion approximation of the Wright-Fisher model assigns probabilities to frequency changes given an N_e_ and the number of generations between sample times. In our model we fixed the within host generation time as either 6 or 12 hours (Geoghegan et al. 2016) and report the findings for the 6 hour generation time below. We then asked what population size makes the observed changes in frequency most likely (Figure 2B). We restricted this analysis to samples taken at least 1 day apart (n = 29), as there was very little change in iSNV frequency in populations sampled twice on the same day (R^2^ = 0.986, Figure 2B and Supplementary Figure 6). The concordance of same day samples suggests that our sampling procedure is reproducible and that less than a generation had passed between samplings. Maximum likelihood optimization of this diffusion model revealed a within host effective population size of 35 (95% CI 26-46, Table 2).

**Figure 2.**
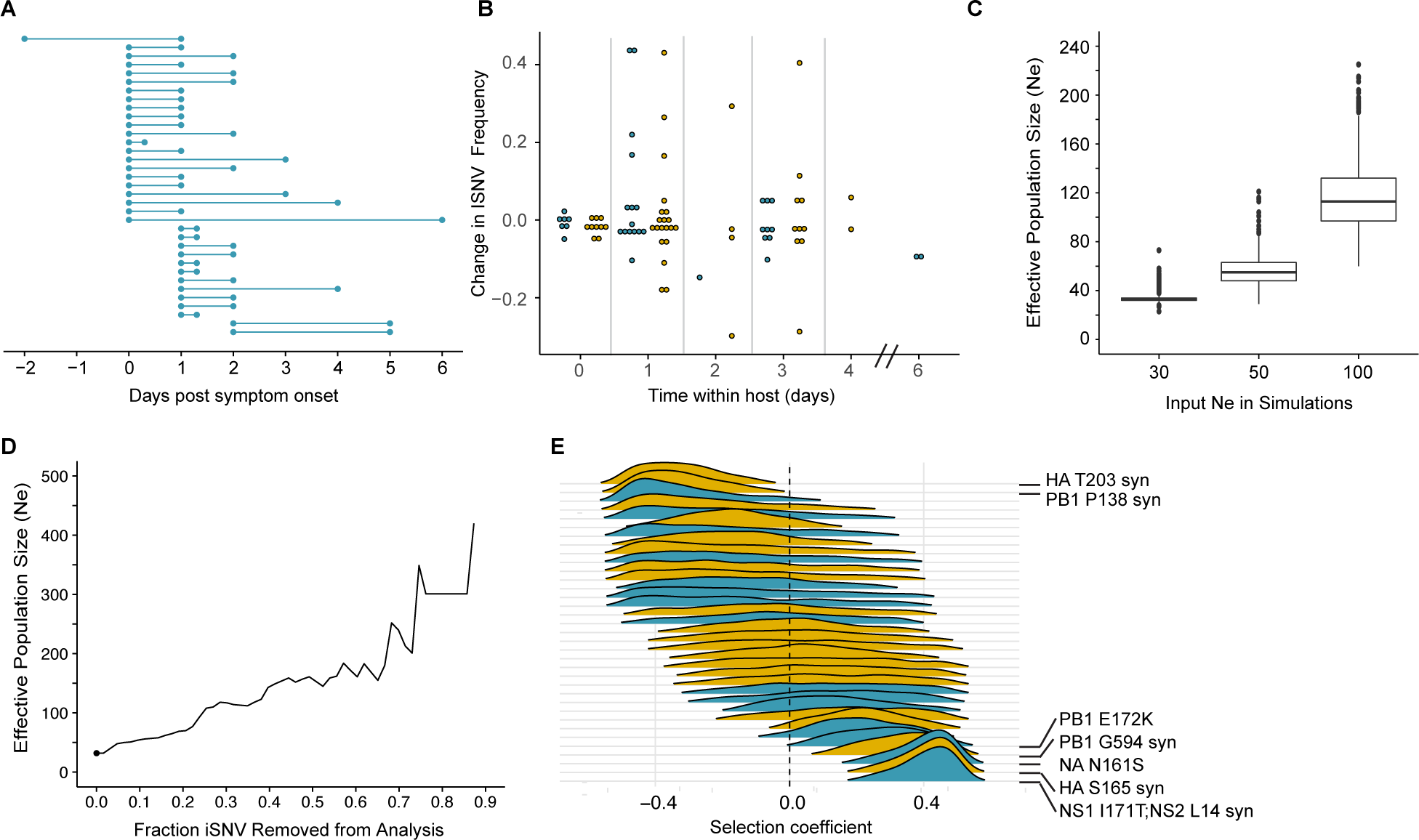
Within host dynamics of IAV. (A) Timing of sample collection for 35 paired longitudinal samples relative to day of symptom onset. Of the 49 total, 35 pairs had minor iSNV present in the first sample. (B) The change in frequency over time for minority nonsynonymous (blue) and synonymous (gold) iSNV identified for the paired samples in (A). (C) The distribution of effective population sizes estimated from 1,000 simulated populations. Simulations were run on populations with characteristics similar to the actual patient-derived populations and with the specified effective population size (x-axis). (D) The effect of iteratively removing iSNV with the most extreme change in frequency (fraction of iSNV removed, x-axis) on the estimated effective population size. The point represents the estimate when all iSNV are included. (E) The posterior distributions of selection coefficients estimated for the 35 iSNV present in isolates sampled one day apart. Distributions are colored according to class relative to the sample consensus sequence, nonsynonymous (blue) synonymous (gold). Variants for which the 95% highest posterior density intervals exclude 0.0 are noted in the margin.

**Table 2.** Within host effective population size of IAV.

| Model | SNV Used | Generation Time (h) | Effective Population Size (95% CI) |
| --- | --- | --- | --- |
| Diffusion approximation | All | 6 | 35 (26-46) |
|  | All | 12 | 17 (13-23) |
| Discrete model | All | 6 | 32 (28-41) |
|  | Nonsynonymous | 6 | 30 (21-40) |
|  | Synonymous | 6 | 37 (27-54) |
|  | All | 12 | 23 (23-29) |
|  | Nonsynonymous | 12 | 19 (19-21) |
|  | Synonymous | 12 | 27 (22-33) |

The diffusion approximation makes several simplifying assumptions, which if violated could influence our findings. To ensure our results were robust to the assumption of a large population, we employed a discrete interpretation of the Wright-Fisher model which makes no assumptions about population size (Williamson & Slatkin 1999). In this case we found an effective population size of 32 (95% CI 28-41), very close to our original estimate (Table 2). Both models assume complete independence of iSNV. To ensure this assumption did not affect our results, we fit the discrete model 1000 times, each time randomly subsetting our data such that only one iSNV per individual was included. This simulates a situation in which all modeled iSNV are independent and our assumption is met. Under these conditions we found a median effective population size of 33 (IQR 32-40), demonstrating negligible bias in the initial analysis due to correlation between iSNV.

As above, most iSNV in the longitudinal samples were rare (< 10%) and many became extinct between samplings. To ensure that our models were capable of accurately estimating the effective population size from such data, we simulated 1000 Wright-Fisher populations with iSNV present at approximately the same starting frequencies as in our data set and an N_e_ of 30, 50, or 100. In these simulations, we found mean N_e_ of 34, 56 and 117 (Figure 2C). These simulations suggest that although this method may slightly overestimate the N_e_, our results are not constrained by the data structure.

To this point, we have assumed that neutral processes are responsible for the observed changes in iSNV frequency within hosts. Although this assumption seems justified at least in part by the analysis above, we tested the robustness of our models by fitting the nonsynonymous (n = 27) and synonymous iSNV (n = 36) separately. Here, we estimated an effective population size of 30 using the nonsynonymous iSNV and an effective population size of 37 using the synonymous iSNV (Table 2). These estimates are very close to those derived from the whole dataset and suggest that nonsynonymous and synonymous mutations are influenced by similar within host processes. To further ensure that our results were not driven by a few outliers subject to strong selection, we ranked iSNV by their change in frequency over time and consecutively removed iSNV with the most extreme changes. We estimated the effective population size at each iteration and found that removing the top 50% most extreme iSNV only increased the effective population size to 161 (Figure 2D). Therefore, our estimates are robust to a reasonable number of non-neutral sites. Finally, we also applied a separate Approximate Bayesian Computational (ABC) method, which uses a non-biased moment estimator in conjunction with ABC to estimate the effective population size of a population as well as selection coefficients for the iSNV present (Foll et al. 2014). This distinct approach relaxes the previous assumption regarding neutrality. We applied this analysis to the 16 longitudinal pairs that were sampled 1 day apart and estimated an effective population of 69. We were unable to reject neutrality for just 7 of the 35 iSNV in this data set (Figure 2E). These seven mutations consisted of 3 nonsynonymous and 4 synonymous mutations and were split between two individuals. None were putative antigenic variants.

### Identification of forty-three transmission pairs

We analyzed virus populations from 85 households with concurrent infections to quantify the level of shared viral diversity and to estimate the size of the IAV transmission bottleneck (Table 1). Because epidemiological linkage does not guarantee that concurrent cases constitute a transmission pair (Petrie et al. 2017), we used a stringent rubric to eliminate individuals in a household with co-incident community acquisition of distinct viruses. We considered all individuals in a household with symptom onset within a 7-day window to be epidemiologically linked. The donor in each putative pair was defined as the individual with the earlier onset of symptoms. We discarded a transmission event if there were multiple possible donors with the same day of symptom onset. Donor and recipients were not allowed to have symptom onset on the same day, unless the individuals were both index cases for the household. In these 6 instances, we analyzed the data for both possible donor-recipient directionalities. Based on these criteria, our cohort had 124 putative household transmission events over 5 seasons (Table 1). Of these, 52 pairs had samples of sufficient quality for reliable identification of iSNV from both individuals.

We next used sequence data to determine which of these 52 epidemiologically linked pairs represented true household transmission events as opposed to coincident community-acquired infections. We measured the genetic distance between influenza populations from each household pair by L1-norm and compared these distances to those of randomly assigned community pairs within each season (Figure 3A, see also trees in Supplementary Figures 2 and 3). While the L1-norm of a pair captures differences between the populations at all levels, in our cohort, it was largely driven by differences at the consensus level. We only considered individuals to be a true transmission pair if they had a genetic distance below the 5th percentile of the community distribution of randomly assigned pairs (Figure 3A). Forty-seven household transmission events met this criterion (Figure 3B). Among these 47 sequence-validated transmission pairs, 3 had no iSNV in the donor and 1 additional donor appeared to have a mixed infection. These four transmission events were removed from our bottleneck analysis as donors without iSNV are uninformative and mixed infections violate model assumptions of site independence (see Methods). We estimated the transmission bottleneck in the remaining 43 high-quality pairs (37 H3N2, 6 H1N1, Figure 3B).

**Figure 3.**
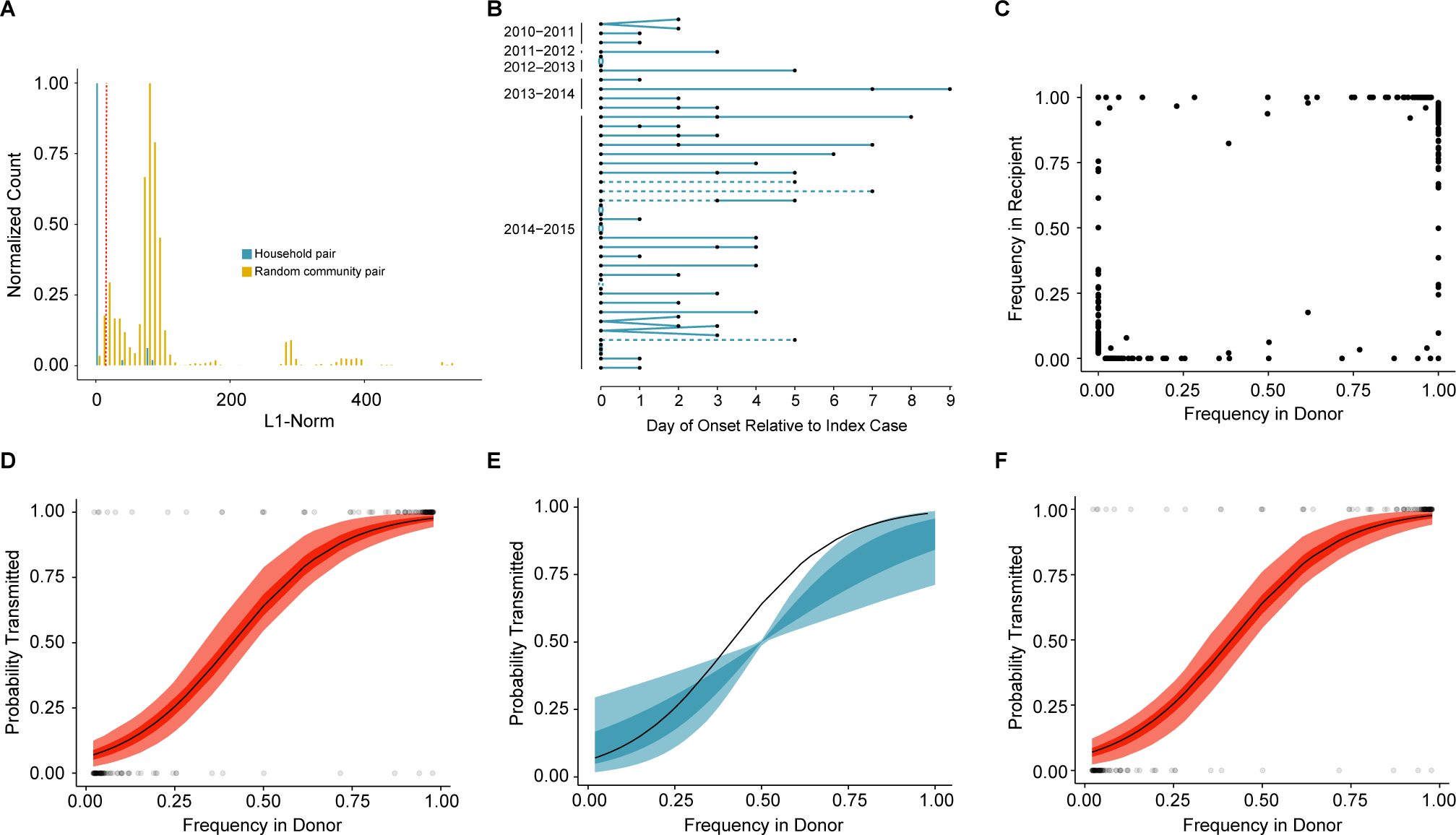
Between host dynamics of IAV. (A) The distribution of pairwise L1-norm distances for household (blue) and randomly-assigned community (gold) pairs. The bar heights are normalized to the height of the highest bar for each given subset (47 for household, 1,592 for community). The red line represents the 5th percentile of the community distribution. (B) Timing of symptom onset for 52 epidemiologically linked transmission pairs. Day of symptom onset for both donor and recipient individuals is indicated by black dots. Dashed lines represent pairs that were removed due to abnormally high genetic distance between isolates, see (A). (C) The frequency of donor iSNV in both donor and recipient samples. Frequencies below 2% and above 98% were set to 0% and 100% respectively. (D) The presence-absence model fit compared with the observed data. The x-axis represents the frequency of donor iSNV with transmitted iSNV plotted along the top and nontransmitted iSNV plotted along the bottom. The black line indicates the probability of transmission for a given iSNV frequency as determined by logistic regression. Similar fits were calculated for 1,000 simulations with a mean bottleneck size of 1.66. Fifty percent of simulated outcomes lie in the darkly shaded region and 95% lie in the lightly shaded regions. (E) The outcome from 1,000 simulated “transmission” events with randomly assigned pairings. The black line represents the observed data, as in (D) the shaded regions represent the middle 50% and 95% of simulated outcomes. The results from the simulated logit models were smoothed by plotting the predicted probability of transmission at 0.02 intervals. (F) The beta-binomial model fit. Similar to (D) except the simulated outcomes are the based on a beta-binomial model using a mean bottleneck of 1.73.

**Figure 4.**
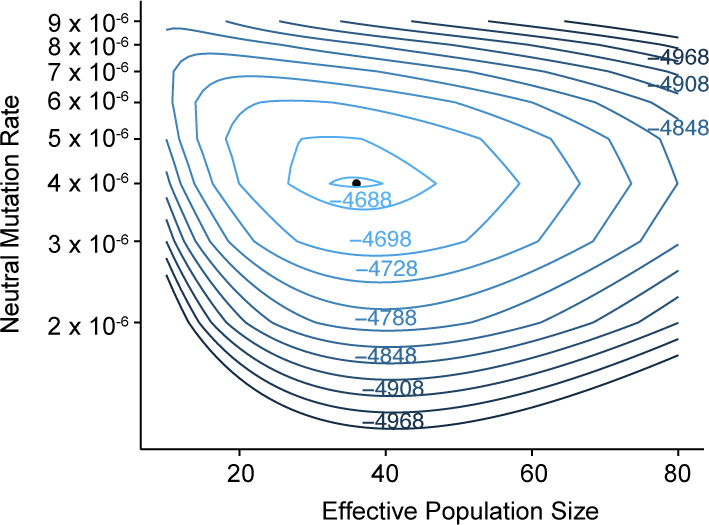
Combined estimates of within host mutation rate and effective population size. Contour plot shows the log likelihood surface for estimates of the effective population size and neutral mutation rate. The point represents the peak (μ = 4x10^−6^, *N_e_* = 36, log likelihood = −4,687). Log likelihoods for each contour are indicated.

A transmission bottleneck restricts the amount of genetic diversity that is shared by both members of a pair. We found that few minority iSNV where polymorphic in both the donor and recipient populations (Figure 3C). Minority iSNV in the donor were either absent or fixed in the recipient (top and bottom of plot). The lack of shared polymorphic sites (which would lie in the middle of the plot in Figure 3C) suggests a stringent effective bottleneck in which only one allele is passed from donor to recipient.

### Estimation of the transmission bottleneck

We applied a simple presence-absence model to quantify the effective transmission bottleneck in our cohort. The presence-absence model considers only whether or not a donor allele is present or absent in the recipient sample. Under this model, transmission is a neutral, random sampling process, and the probability of transmission is simply the probability that the iSNV will be included at least once in the sample given its frequency in the donor and the sample size, or bottleneck. We estimated a distinct bottleneck for each transmission pair and assumed these bottlenecks followed a zero-truncated Poisson distribution. This model also assumes that the sensitivity for detection of transmitted iSNVs is perfect and that each genomic site is independent of all others. We then used maximum likelihood optimization to determine the distribution of bottleneck sizes that best fit the data. We found a zero-truncated Poisson distribution with a mean of 1.66 (lambda = 1.12; 0.51-1.99, 95% CI) best described the data. This distribution indicates that the majority of bottlenecks are 1, and that very few are greater than 5 (probability 0.2%). There were no apparent differences between H3N2 and H1N1 pairs. The model fit was evaluated by simulating each transmission event 1,000 times. The presence or absence of each iSNV in the recipient was noted and the probability of transmission given donor frequency determined. The range of simulated outcomes matched the data well, which suggests that transmission is a selectively neutral event characterized by a stringent bottleneck (Figure 3D).

The majority of transmitted iSNV were fixed in the recipients. Although this trend matches the expectation given a small bottleneck, these data could also be consistent with a model in which the probability of transmission is determined by the frequency at which iSNV are found at the community level. To ensure our bottleneck estimates were an outcome of neutral transmission and not an artifact of the larger community population structure or selection for the community consensus, we created a null model by randomly assigning community “recipient-donor” pairings. Each community “recipient” was drawn from the pool of individuals that were infected after the “donor” but in the same season and with the same subtype as the donor. We then identified whether or not each donor iSNV was found in the community recipient and determined the relationship between “donor” frequency and probability of “transmission” for 1,000 such simulations. Given the low level of diversity in our cohort, we predicted that rare iSNV would be unlikely to be found in a random sample, while the major alleles should be fixed in most random pairs. This trend is clearly demonstrated in Figure 3E. It is also clear that this null model fit the data much more poorly than the presence/absence model, suggesting that the observed data in our bona fide transmission pairs were not a product of community metapopulation structure, but rather an outcome of neutral sampling events.

Because our bottleneck estimates were much lower than what has previously been reported for human influenza (Poon et al. 2016), we investigated the impact that our simplifying assumptions could have on our results. In particular, the presence-absence model assumes perfect detection of variants in donor and recipient, and it can therefore underestimate the size of a bottleneck in the setting of donor-derived variants that are transmitted but not detected in the recipient. These “false negative” variants can occur when the frequency of an iSNV drifts below the level of detection (e.g. 2% frequency) or when the sensitivity of sequencing is less than perfect for variants at that threshold (e.g. 15% sensitivity for variants at a frequency 2-5%). To determine the impact of sequencing sensitivity and specificity on our bottleneck estimates, we re-called variants using our original pipeline without the 2% frequency cut-off. As shown in Supplementary Table 1, this increases the sensitivity of iSNV detection in the 1-5% frequency range, and also the number of false positive variant calls (McCrone & Lauring 2016). This analysis only slightly increased average transmission bottleneck to 2.10 (lambda = 1.67; 0.91-2.71, 95% CI), and indicates that our results are not biased by the added stringency used in the initial analysis (Supplementary Figures 7A and 7B).

To further investigate the impact of sequencing accuracy on our estimates, we inferred minor variants in our current pipeline (see above and methods) without a frequency cutoff. Ultimately, this reduced variant calling to a count method at a number of positions and greatly increased the number of shared minority iSNV in our samples (Supplementary Figure 7C). Many of these presumed false positive variant calls were at similar frequencies (0.1-2%) in donor and recipient. As such, the “apparent transmission” of rare variants drives an inflated estimate of the transmission bottleneck (118, see Supplementary Figure 7D). Simulation showed that this inflated bottleneck no longer fit the trend in the data, likely because the model is now forced to accommodate shared iSNV that are biased toward sequencing error as opposed to the actual transmission process.

We also estimated bottleneck size using a beta binomial model, which Leonard *et al.* have used to account for the stochastic loss of transmitted variants. This model allows for a limited amount of time-independent genetic drift within the recipient (Sobel Leonard et al. 2017), and we modified it to also account for our benchmarked sensitivity for rare variants (Supplementary Table 1, Current Pipeline). For all donor-derived iSNV that were absent in the recipient, we estimated the likelihood that these variants were transmitted but either drifted below our level of detection or drifted below 10% and were missed by our variant identification. Despite the relaxed assumptions provided by this modified beta binomial model, maximum likelihood estimation only marginally increased the average bottleneck size (mean 1.73: lambda 1.22; 0.57-2.17, 95%CI) relative to the simpler presence-absence model. We simulated transmission and subsequent random drift using the beta binomial model and the estimated bottleneck distribution as above (Figure 3F). Although the model matched the data well, the fit was not better than that of the presence-absence model (AIC 83.0 for beta-binomial compared to 76.7 for the presence-absence model).

### The mutation rate of influenza A virus within human hosts

The stringent influenza transmission bottleneck suggests that most infections are founded by one lineage and develop under essentially clonal processes. The diffusion approximation to the Wright-Fisher model (see above and Figure 2) can be used to predict the rate at which homogenous populations diversify from a clonal ancestor as a function of mutation rate and effective population size (Rouzine et al. 2001). Maximum likelihood optimization of this model suggested an *in vivo* neutral mutation rate of 4×10^−6^ mutations per nucleotide per replication cycle and a within host effective population size of 36 (given a generation time of 6 hours). These estimates are consistent with those above (Table 2). As we have recently estimated that 13% of mutations in influenza A virus are neutral (Visher et al. 2016), we estimated that the true *in vivo* mutation rate would be approximately 8 fold higher than our neutral rate – on the order of 3-4 × 10^−5^. This in vivo mutation rate is close to our recently published estimate of influenza A mutation rates in epithelial cells by fluctuation test (Pauly et al. 2017) and within the range of other estimates for IAV (Sanjuán et al. 2010).

## Discussion

We find that seasonal influenza A viruses replicate within and spread among human hosts with very small effective population sizes. Because we used viruses collected over five influenza seasons from individuals enrolled in a prospective household cohort, these dynamics are likely to be broadly representative of many seasonal influenza infections in their natural transmission context. Our results are further strengthened by the use of a validated sequence analysis pipeline and models that are robust to the underlying assumptions. The small effective size of intrahost populations and the tight effective transmission bottleneck suggest that stochastic processes, such as genetic drift, dominate influenza virus evolution at the level of individual hosts. This stands in contrast to prominent role of positive selection in the global evolution of seasonal influenza.

While influenza virus populations are subject to continuous natural selection, selection is an inefficient driver of evolution in small populations (Rouzine et al. 2001). Despite a large viral copy number, our findings demonstrate that intrahost populations of influenza behave like much smaller populations. We therefore expect stochastic fluctuations to be the major force driving the fixation of novel variants within human hosts. This finding contradicts previous studies, which have found signatures of adaptive evolution in infected hosts (Gubareva et al. 2001; Rogers et al. 2015; Ghedin et al. 2011; Sobel Leonard et al. 2016). However, these studies rely on data from infections in which selective pressures are likely to be particularly strong (e.g. due to drug treatment or infection with a poorly adapted virus), or in which the virus has been allowed to propagate for extended periods of time (Xue et al. 2017). Under these conditions, one can identify the action of positive selection on within host populations. We suggest that these are important and informative exceptions to the drift regime defined here.

We used both a simple presence-absence model and a more complex beta binomial model to estimate an extremely tight transmission bottleneck. The estimation of a small bottleneck size is driven by low within-host diversity and very few minority iSNV shared among individuals in a transmission pair. While our methods for variant calling may be more conservative than those used in similar studies, we found that relaxing our variant calling criteria led to the inclusion of false positive variants that inflated our estimates. Furthermore, the beta binomial model accounts for false negative iSNV (i.e. variants that are transmitted but not detected in the donor), which can lead to underestimated transmission bottlenecks (Sobel Leonard et al. 2017). Our formulation of this model incorporates empirically determined sensitivity and specificity metrics to account for both false negative iSNV and false positive iSNV (McCrone & Lauring 2016). Finally, if rare, undetected, iSNV were shared between linked individuals, we would expect to see transmission of more common iSNV (frequency 5-10%), which we can detect with high sensitivity. In our data, the transmission probability iSNVs > 5% frequency in the donor were also well predicted by small bottleneck size (Figure 3D).

Although the size of our transmission bottleneck is consistent with estimates obtained for other viruses and in experimental animal models of influenza (Zwart & Elena 2015; Varble et al. 2014), it differs substantially from the only other study of bottlenecks in natural human infection (Poon et al. 2016; Sobel Leonard et al. 2017). While there are significant differences in the design and demographics of the cohorts, the influenza seasons under study, and sequencing methodology (Kugelman et al. 2017), the bottleneck size estimates are fundamentally driven by the amount of viral diversity shared among individuals in a household. Importantly, we used both epidemiologic linkage and the genetic relatedness of viruses in households to define transmission pairs and to exclude confounding from the observed background diversity in the community. We find that household transmission pairs and randomly assigned community pairs had distinct patterns of shared consensus and minority variant diversity. The comparison to random community pairs is important, as an unexplained aspect of the work of Poon et al. is that rare iSNV were frequently shared by randomly selected individuals, and more common ones were not (Poon et al. 2016).

Our estimates of IAV population dynamics are consistent across three separate models and partitions of the data. The measurements of shared diversity are influenced by both between and within host processes, and the transmission bottleneck is entirely consistent with the small within host population size derived from the longitudinal samples. We also jointly estimated the *in vivo* mutation rate and effective population size based on the frequency distribution of minor alleles observed in the entire cohort. This model assumed a small transmission bottleneck, produced a mutation rate that this consistent with previous estimates, and independently reproduced the within host population size estimate. Given the concordance among these distinct approaches, it is unlikely that our findings are biased by hidden assumptions or model limitations.

Accurately modeling and predicting influenza virus evolution requires a thorough understanding of the virus' population structure. Some models have assumed a large intrahost population and a relatively loose transmission bottleneck (Geoghegan et al. 2016; Russell et al. 2012; Peck et al. 2015). Here, adaptive iSNV can rapidly rise in frequency and low frequency variants can have a high probability of transmission. In such a model, it would be possible for the highly pathogenic H5N1 virus to develop the requisite 4-5 mutations to become transmissible through aerosols during a single acute infection of a human host (Herfst et al. 2012; Russell et al. 2012). Although the dynamics of emergent avian influenza and human adapted seasonal viruses likely differ (Petrova & Russell 2017), our work suggests that fixation of multiple mutations over the course of a single acute infection is unlikely.

While it may seem counterintuitive that influenza evolution is dominated by drift on local scales and positive selection on global scales, these models are certainly not in conflict. Within individuals we have shown that the effective population is quite small, which suggests that selection is inefficient. Indeed, we have deeply sequenced 332 intrahost populations from 283 individuals collected over more than 11,000 person-seasons of observation and only identified a handful of minority antigenic variants with little evidence for positive selection (this work and (Debbink et al. 2017)). However, with several million infected individuals each year, even inefficient processes and rare events are likely to happen at a reasonable frequency on a global scale.

## Methods

### Description of the cohort

The HIVE cohort (Monto et al. 2014; Ohmit et al. 2013; Ohmit et al. 2015; Ohmit et al. 2016; Petrie et al. 2013), established at the UM School of Public Health in 2010, enrolled and followed households of at least 3 individuals with at least two children <18 years of age; households were then followed prospectively throughout the year for ascertainment of acute respiratory illnesses. Study participants were queried weekly about the onset of illnesses meeting our standard case definition (two or more of: cough, fever/feverishness, nasal congestion, sore throat, body aches, chills, headache if ≥3 yrs old; cough, fever/feverishness, nasal congestion/runny nose, trouble breathing, fussiness/irritability, decreased appetite, fatigue in <3 yrs old), and the symptomatic participants then attended a study visit at the research clinic on site at UM School of Public Health for sample collection. For the 2010-2011 through 2013-2014 seasons, a combined nasal and throat swab (or nasal swab only in children < 3 years of age) was collected at the onsite research clinic by the study team. Beginning with the 2014-2015 seasons, respiratory samples were collected at two time points in each participant meeting the case definition; the first collection was a self‐ or parent-collected nasal swab collected at illness onset. Subsequently, a combined nasal and throat swab (or nasal swab only in children < 3 years of age) was collected at the onsite research clinic by the study team. Families with very young children (< 3 years of age) were followed using home visits by a trained medical assistant.

Active illness surveillance and sample collection for cases were conducted October through May and fully captured the influenza season in Southeast Michigan in each of the study years. Data on participant, family and household characteristics, and on high-risk conditions were additionally collected by annual interview and review of each participant's electronic medical record. In the current cohort, serum specimens were also collected twice yearly during fall (November-December) and spring (May-June) for serologic testing for antibodies against influenza.

This study was approved by the Institutional Review Board of the University of Michigan Medical School, and all human subjects provided informed consent.

### Identification of influenza virus

Respiratory specimens were processed daily to determine laboratory-confirmed influenza infection. Viral RNA was extracted (Qiagen QIAamp Viral RNA Mini Kit) and tested by RT-PCR for universal detection of influenza A and B. Samples with positive results by the universal assay were then subtyped to determine A(H3N2), A(H1N1), A(pH1N1) subtypes and B(Yamagata) and B(Victoria) lineages. We used primers, probes and amplification parameters developed by the Centers for Disease Control and Prevention Influenza Division for use on the ABI 7500 Fast Real-Time PCR System platform. An RNAseP detection step was run for each specimen to confirm specimen quality and successful RNA extraction.

### Quantification of viral load

Quantitative reverse transcription polymerase chain reaction (RT-qPCR) was performed on 5μl RNA from each sample using CDC RT-PCR primers InfA Forward, InfA Reverse, and InfA probe, which bind to a portion of the influenza M gene (CDC protocol, 28 April 2009). Each reaction contained 5.4μl nuclease-free water, 0.5μl each primer/probe, 0.5μl SuperScript III RT/Platinum Taq mix (Invitrogen 111732) 12.5μl PCR Master Mix, 0.1μl ROX, 5μl RNA. The PCR master mix was thawed and stored at 4°C, 24 hours before reaction set-up. A standard curve relating copy number to Ct value was generated based on 10-fold dilutions of a control plasmid run in duplicate.

### Illumina library preparation and sequencing

We amplified cDNA corresponding to all 8 genomic segments from 5μSuperScript III One-Step RT-PCR Platinum Taq HiFi Kit (Invitrogen 12574). Reactions consisted of 0.5μl Superscript III Platinum Taq Mix, 12.5μl 2x reaction buffer, 6μl DEPC water, and 0.2μl of 10μM Uni12/Inf1, 0.3μl of 10μM Uni12/Inf3, and 0.5μl of 10μM Uni13/Inf1 universal influenza A primers (Zhou et al. 2009). The thermocycler protocol was: 42°C for 60 min then 94°C for 2 min then 5 cycles of 94°C for 30 sec, 44°C for 30 sec, 68°C for 3 min, then 28 cycles of 94°C for 30 sec, 57°C for 30 sec, 68°C for 3 min. Amplification of all 8 segments was confirmed by gel electrophoresis, and 750ng of each cDNA mixture were sheared to an average size of 300 to 400bp using a Covaris S220 focused ultrasonicator. Sequencing libraries were prepared using the NEBNext Ultra DNA library prep kit (NEB E7370L), Agencourt AMPure XP beads (Beckman Coulter A63881), and NEBNext multiplex oligonucleotides for Illumina (NEB E7600S). The final concentration of each barcoded library was determined by Quanti PicoGreen dsDNA quantification (ThermoFisher Scientific), and equal nanomolar concentrations were pooled. Residual primer dimers were removed by gel isolation of a 300-500bp band, which was purified using a GeneJet Gel Extraction Kit (ThermoFisher Scientific). Purified library pools were sequenced on an Illumina HiSeq 2500 with 2x125 nucleotide paired end reads. All raw sequence data have been deposited at the NCBI sequence read archive (BioProject submission ID: SUB2951236). PCR amplicons derived from an equimolar mixture of eight clonal plasmids, each containing a genomic segment of the circulating strain were processed in similar fashion and sequenced on the same HiSeq flow cell as the appropriate patient derived samples. These clonally derived samples served as internal controls to improve the accuracy of variant identification and control for batch effects that confound sequencing experiments.

### Identification of iSNV

Intrahost single nucleotide variants were identified in samples that had greater than 10^3^ genomes/μl and an average coverage >1000x across the genome. Variants were identified using DeepSNV and scripts available at https://github.com/lauringlab/variant_pipeline as described previously (McCrone & Lauring 2016) with a few minor and necessary modifications. Briefly, reads were aligned to the reference sequence (H3N2 2010-2011 & 2011-2012: GenBank CY121496-503, H3N2 2012-2013:GenBank KJ942680-8, H3N2 2014-2015: Genbank CY207731-8, H1N1 GenBank: CY121680-8) using Bowtie2 (35). Duplicate reads were then marked and removed using Picard (http://broadinstitute.github.io/picard/). We identified putative iSNV using DeepSNV. Bases with phred <30 were masked. Minority iSNV (frequency <50%) were then filtered for quality using our empirically determined quality thresholds (p-value <0.01 DeepSNV, average mapping quality >30, average Phred >35, average read position between 31 and 94). To control for PCR errors in samples with lower input titers, all isolates with titers between 10^3^ and 10^5^ genomes/μl were processed and sequenced in duplicate. Only iSNV that were found in both replicates were included in down stream analysis. The frequency of the variant in the replicate with higher coverage at the iSNV location was assigned as the frequency of the iSNV. Finally, any SNV with a frequency below 2% was discarded.

Given the diversity of the circulating strain in a given season, there were a number of cases in which isolates contained mutations that were essentially fixed (>95%) relative to the plasmid control. Often in these cases, the minor allele in the sample matched the major allele in the plasmid control. We were, therefore, unable to use DeepSNV in estimating the base specific error rate at this site for these minor alleles and required an alternative means of eliminating true and false minority iSNV. To this end we applied stringent quality thresholds to these putative iSNV and implemented a 2% frequency threshold. In order to ensure we were not introducing a large number of false positive iSNV into our analysis, we performed the following experiment. Perth (H3N2) samples were sequenced on the same flow cell as both the Perth and Victoria (H3N2) plasmid controls. Minority iSNV were identified using both plasmid controls. This allowed us to identify rare iSNV at positions in which the plasmid controls differed both with and without the error rates provided by DeepSNV. We found that at a frequency threshold of 2% the methods were nearly identical (NPV of 1, and PPV of 0.94 compared to DeepSNV).

### Overview of models for effective population size

We estimated the effective population size using two separate interpretations of a Wright-Fisher population (Ewens 2004). At its base, the Wright-Fisher model describes the expected changes in allele frequency of an ideal population, which is characterized by non-overlapping generations, no migration, no novel mutation, and no population structure. We then asked what size effective population would make the changes in frequency observed in our dataset most likely. We calculated these values using two applications of the Wright-Fisher model (i) a diffusion approximation (Kimura 1955)and (ii) a maximum likelihood approach based on the discrete interpretation (Williamson & Slatkin 1999).

For these estimates we restricted our analysis to longitudinal samples from a single individual that were separated by at least 1 day and only used sites that were polymorphic in the initial sample (29 of the 49 total serial sample pairs). We modeled only the iSNV that were the minor allele at the first time point, and we assumed a within host generation time of either 6 or 12 hours as proposed by Geoghegan *et al.* (Geoghegan et al. 2016).

### Diffusion approximation

The diffusion approximation was first solved by Kimura in 1955 (Kimura 1955). This approximation to the discrete Wright-Fisher model has enjoyed widespread use in population genetics as it allows one to treat the random time dependent probability distribution of final allele frequencies as a continuous function (e.g. (Zanini et al. 2017; Kimura & Ohta 1969; Kimura 1971; Myers et al. 2008)). Here, we also included the limitations in our sensitivity to detect rare iSNV by integrating over regions of this probability density that were either below our limit of detection or within ranges where we expect less than perfect sensitivity as follows.

Let *P*(*p_0_,p_t_,t|N_e_*)be the time dependent probability of a variant drifting from an initial frequency of *p_0_* at time 0*p_t_* at time *t* generations given an effective population size of *N_e_* where 0<*p_t_*<1.

The time dependent derivative of this probability has been defined using the Kolmogorov forward equation (Kimura 1955) and for haploid populations is:

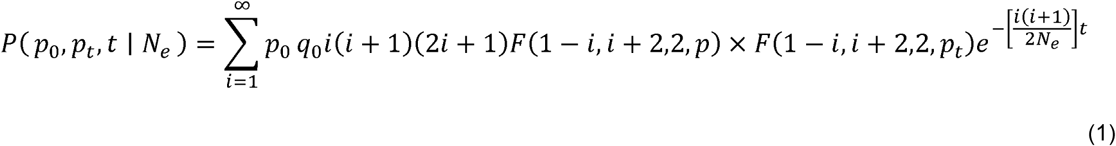

Where *q* = 1 – *p* and *F* is the hypergeometric function. We approximated the infinite sum by summing over the first 50 terms. When we added an additional 50 terms (100 in total) we found no appreciable change in the final log likelihoods.

We denote the frequency of allele that is not observed at the second time point as *p_t_* ≈ 0 and the probability of such an event as *P*(*p_0_,p_t_ ≈ 0,t| N_e_*). This probability is given in equation 2 as the sum of the probability that the variant is truly lost by generation (i.e. the other allele is fixed *P*(*q_0_,q_t_ = 1,t|N_e_*), the probability that it is present but below the limit of detection (i.e. *P*(*p_0_,p_t_ ≈ 0,t| 0<*p_t_*<0.02 N_e_*)) and the probability the variant is not detected due to low sensitivity for rare variant detection (i.e. *P*(*p_0_,p_t_ ≈ 0,t| 0.02<*p_t_*<0.1 N_e_*)). The probability of not observing an allele at the second time is then

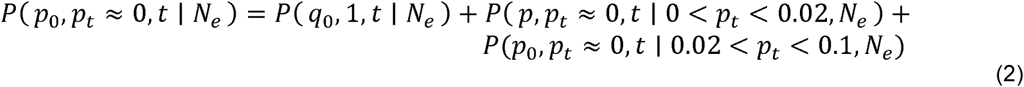

The first term in equation 2 is adapted from Kimura, 1955 as

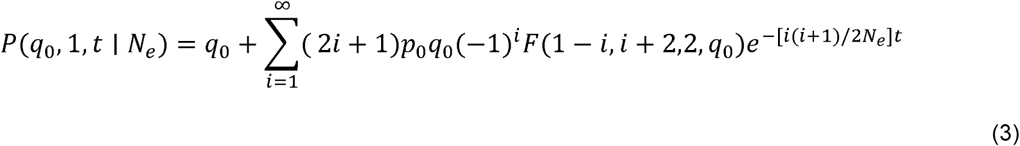

Where q is defined as above. (Note that this is simply the probability of fixation for a variant at initial frequency q). As in equation 1 the infinite sum was approximated with a partial sum of 50 terms.

The probability of the allele drifting below our limit of detection can be found by integrating equation 1 between 0 and our limit of detection, 0.02. This was done numerically using the python package scipy.

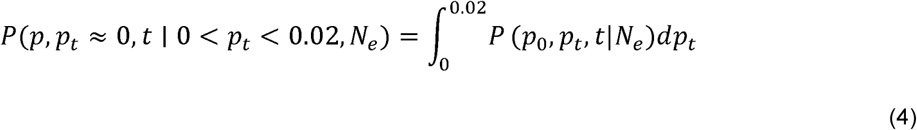

Finally, the probability of an iSNV being present at the second time point, but escaping detection, is given by the integral of equation 1 between our benchmarked frequencies (0.02,0.05) times the false negative rate for that range. Here, we assumed the entire range had the same sensitivity as the benchmarked frequency at the lower bound and rounded recipient titers down
to the nearest log_10_ titer (e.g. 10^3^,10^1^, 10^5^). We also assumed perfect sensitivity above 10%.

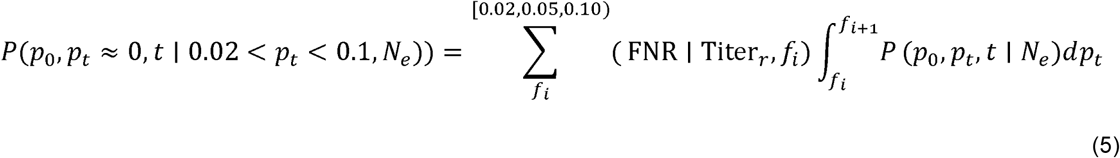

Where (FNR | Titer_*r*_,*f_i_*) is the false negative rate given the frequency and the sample titer (See Supplemental Table 1) and *P*(*p_0_,p_t_,t| N_e_*) is defined in equation 1.

The log likelihood of an effective population size is the sum of the log of *P*(*p_0_,p_t_,t| N_e_*) for each minor allele in the data set, where either the position is polymorphic at time *t* (i.e. equation 1) or the allele is not observed at time *t* (i.e. equation 2).

### Discrete Wright-Fisher estimation of *N_e_*

The diffusion approximation treats changes in frequency as a continuous process because it assumes sufficiently large *N_e_*. That assumption can be relaxed, and the effective population size can be determined, by applying a maximum likelihood method developed by Williamsom and Slatkin 1999 (Williamson & Slatkin 1999). In this model, the true allele frequencies move between discrete states (i.e. the frequency must be of the form *i*/*N_e_* where *i* is a whole number in the range [0,*N_e_*]. In the original application, allele counts were used, and sampling error was added to the model as a binomial distribution with n determined by the sample size. Here, we use the frequencies available from next generation sequencing and estimate sampling error as a normal distribution with mean equal to the observed frequency and a standard deviation equal to that observed in our benchmarking study for the 10^4^ genomes/μl samples (*σ* = 0.014) (McCrone & Lauring 2016).

In this model, the probability of observing an allele frequency shift from *p̂_0_* to *p̂_t_* in generations provided an effective population of *N_e_* is the probability of observing *p̂_0_* given some initial state *p*_0_ and the probability of the population having that state, times the probability of observing *p̂_t_* given some final state *p_t_* and the probability of moving from the initial to the final state summed across all possible states.

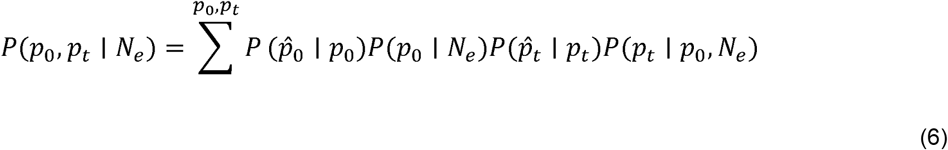

Where *p̂_x_* are the observed probabilities and *p_x_* are the real ones (of the form *i*/*N_e_* discussed above). The likelihood of observing a given frequency *p̂_x_* given a defined state *p_x_* is given by the likelihood of drawing *p̂_x_* from a normal distribution with mean *p_x_* and standard deviation 0.014.

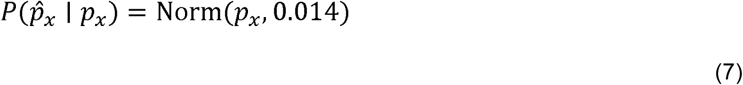

As in Williamson and Slatkin 1999, we assume a uniform prior on the initial state. Because we know that our specificity is near perfect (above 2%, Supplemental Table 1) and we restrict our analysis to only polymorphic sites, the probability of any initial state is given by

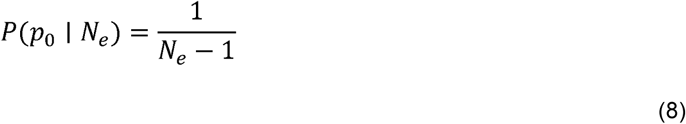

and finally the probability of moving from one state to another in *t* generations is given by

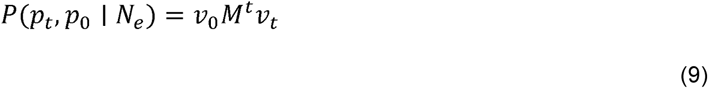

Where M is a square transmission matrix with *N_e_*+1 rows and columns. Where m_i,j_ is the probability of going from the ith configuration to the jth or the probability of drawing *j* – 1 out of binomial distribution with mean (*i* – 1)/*N_e_* and a sample size *N_e_*,*v_0_* is a row vector of initial frequencies *p_0_* with 100% chance of initial state *p_0_*, and *v_t_* is column vector of the frequencies at time point *t* with 100% chance of the final state. In other words *v_0_* is a row vector of *N_e_*+1 states with 0 everywhere except in the ith position where 
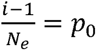
, and *v_t_* is a column vector of *N_e_*+1 states with 0 everywhere except the jth position where 
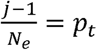
 Using the scalar and cumulative properties of matrix multiplication equation 6 reduces to

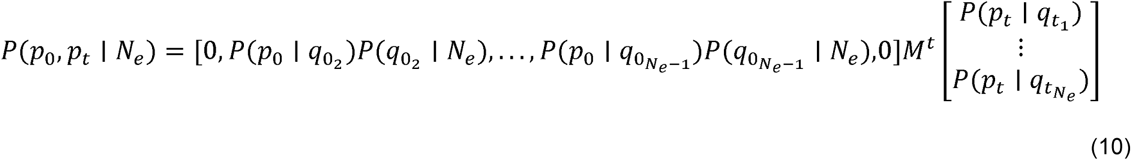

The first and last entries in *v_0_* are 0 because we assume all measured sites represent polymorphisms at the first time of sampling. As above, the log likelihood of a given population size is then simply the sum of the log of *P*(*p̂_0_,p̂_t_,t| N_e_*) for each minor allele in the data set.

### Simulations

To simulate within host evolution we set *N_e_* in equation 10 to either 30, 50 or 100. For each minor allele we used the closest available non-zero state given the effective population size as the starting state. We then calculated the probability of moving to any other state and selected a final state from this distribution. We then drew a final measured frequency from the normal distribution to account for measurement errors.

### ABC model

We estimated both the effective population size and selection coefficients using the approximate Bayesian computation (ABC) described in (Foll et al. 2014) with the WFACB_v1.1 software provided in (Foll et al. 2015). In its current implementation, this analysis requires the same time points for each sample, and we restricted this analysis to longitudinal samples taken 1 day apart. This subset constitutes 16 of the 29 modeled longitudinal samples. Briefly, we subsampled polymorphic sites to 1,000x coverage to estimate allele counts from frequency data as in (Foll et al. 2014). We then estimated the prior distribution of the effective population size using 10,000 bootstrap replicates. We selected a uniform distribution on the range [−0.5,0.5] as the prior distribution for the selection coefficients. The posterior distributions were determined from accepting the top 0.01% of 100,000 simulations.

### Overview of models used for estimating the transmission bottleneck

We model transmission as a simple binomial sampling process (Sobel Leonard et al. 2017). In our first model, we assume any transmitted iSNV, no matter the frequency, will be detected in the recipient. In the second, we relax this assumption and account for false negative iSNV in the recipient. To include the variance in the transmission bottlenecks between pairs we use maximum likelihood optimization to fit the average bottleneck size assuming the distribution follows a zero-truncated Poisson distribution.

### Presence/Absence model

The presence/absence model makes several assumptions. We assume perfect detection of all transmitted iSNV in the recipient. For each donor iSNV, we measure only whether or not the variant is present in the recipient. Any iSNV that is not found in the recipient is assumed to have not been transmitted. We also assume the probability of transmission is determined only by the frequency of the iSNV in the donor at the time of sampling (regardless of how much time passes between sampling and transmission). The probability of transmission is simply the probability that the iSNV is included at least once in a sample size equal to the bottleneck. Finally, we assume all genomic sites are independent of one another. For this reason, we discarded the one case where the donor was likely infected by two strains, as the iSNV were certainly linked.

In our within host models, we only tracked minor alleles as in our data set we only ever find 2 alleles at each polymorphic site. In this case, the frequency of the major allele is simply one minus the frequency of the minor allele. Because the presence/absence model is unaware of the frequency of alleles in the recipient we must track both alleles at each donor polymorphic site.

Let *A_1_* and *A_2_* be alleles in donor *j* at genomic site *i*. Let *P*(*A_1_*) be the probability that *A_1_* is the only transmitted allele. There are three possible outcomes for each site. Either only *A_1_* is transmitted, only *A_2_* is transmitted, or both *A_1_* and *A_2_* are transmitted. The probability of only *A_1_* being transmitted given a bottleneck size of *N_b_* is

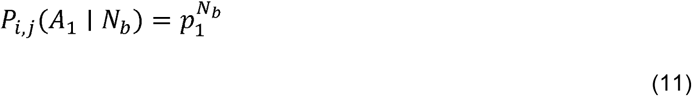

where *p_1_* is the frequency of *A_1_* in the donor. In other words, this is simply the probability of only drawing *A_1_* in *N_b_* draws. The probability that only *A_2_* is transmitted is similarly defined.

The probability of both alleles being transmitted is given by

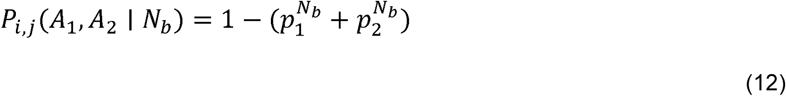

where *p_1_* and *p_2_* are the frequencies of the alleles respectively. This is simply the probability of not picking only *A_1_* or only *A_2_* in *N_b_* draws.

This system could easily be extended to cases where there are more than 2 alleles present at a site; however, that never occurs in our data set.

For ease we will denote the likelihood of observing the data at a polymorphic site *i* in each donor *j* given the bottleneck size *N_b_* as *P_i,j_*(*N_b_*) where *P_i,j_*(*N_b_*) is defined by equation 11 if only one allele is transmitted and equation 12 if two alleles are transmitted.

The log likelihood of a bottleneck of size *N_b_* is given by

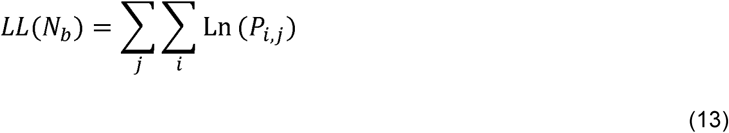

where *i,j* refers to the th polymorphic site in the *i*th donor. This is the log of the probability of observing the data summed over all polymorphic sites across all donors.

Because the bottleneck size is likely to vary across transmission events, we used maximum likelihood to fit the bottleneck distribution as oppose to fitting a single bottleneck value. Under this model we assumed the bottlenecks were distributed according to a zero-truncated Poisson distribution parameterized by *λ*. The likelihood of observing the data given a polymorphic site *i* in donor *j* and *λ* is

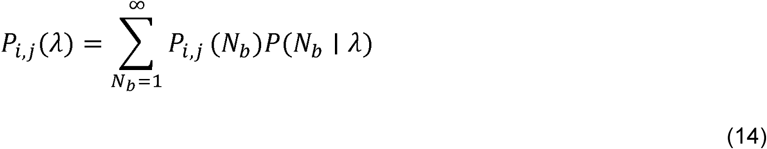

where *P_i,j_*(*N_b_*) is defined as above, *P*(*N_b_* | *λ*)is the probability of drawing a bottleneck of size *N_b_* from a zero-truncated Poisson distribution with a mean of 
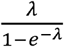
. The sum is across all possible *N_b_* defined on [1,∞]. For practical purposes, we only investigated bottleneck sizes up to 100, as initial analyses suggested *λ* is quite small and the probability of drawing a bottleneck size of 100 from a zero-truncated Poisson distribution with *λ* = 10 is negligible. We follow this convention whenever this sum appears.

The log likelihood of *λ* for the data set is given by

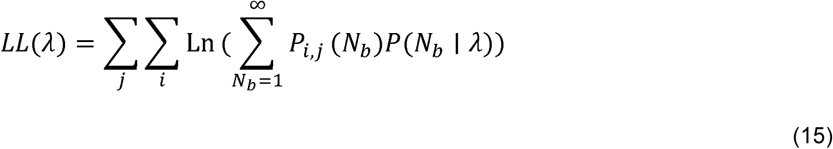

### Beta Binomial model

The Beta binomial model is explained in detail in Leonard *et al.* (Sobel Leonard et al. 2017). It is similar to the presence/absence model in that transmission is modeled as a simple sampling process; however, it relaxes the following assumptions. In this model, the frequencies of transmitted variants are allowed to change between transmission and sampling according a beta distribution. The distribution is not dependent on the amount of time that passes between transmission and sampling, but rather depends on the size of the founding population (here assumed to equal to *N_b_*) and the number of variant genomes present in founding population *k*. Note the frequency in the donor is assumed to be the same between sampling and transmission.

The equations below are very similar to those presented by Leonard *et al.* with one exception. Because we know the sensitivity of our method to detect rare variants based on the expected frequency and the titer, we can include the possibility that iSNV are transmitted but are missed due to poor sensitivity. Because the beta binomial model is aware of the frequency of the iSNV in the recipient, no information is added by tracking both alleles at a genomic site *i*.

Let *p_i,j_d__* represent the frequency of the minor allele frequency at position *i* in the donor of some transmission pair *j*. Similarly, let *p_i,j_r__* be the frequency of that same allele in the recipient of the *j*th transmission pair. Then, as in Leonard *et al.*, the likelihood of some bottleneck *N_b_* for the data at site *i* in pair *j* where the minor allele is transmitted is given by

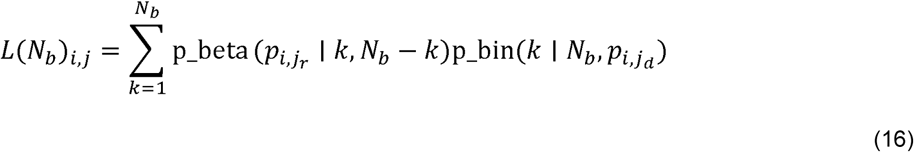

Where p_beta is the probability density function for the beta distribution and p_bin is the probability mass function for the binomial distribution.

This is the probability density that the transmitted allele is found in the recipient at a frequency of *p_i,j_r__* given that the variant was in *k* genomes in a founding population of size *N_b_* times the probability of drawing *k* variant genomes in a sample size of *N_b_* and a variant frequency of *p_i,j_d__*. This is then summed for all possible *k* where 1 ≤ *k* ≤ *N_b_*.

As in equation 14 the likelihood of a zero truncated Poisson with a mean of 
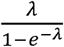
 given this transmitted variants is then given by

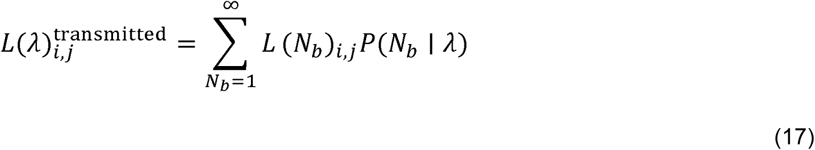

This is simply the likelihood of each *N_b_* weighted by the probability of drawing a bottleneck size of *N_b_* from bottleneck distribution.

In this model, there are three possible mechanisms for a donor iSNV to not be detected in the recipient. (i) The variant was not transmitted. (ii) The variant was transmitted but is present below our level of detection (2%). (iii) The variant was transmitted and present above our level of detection but represents a false negative in iSNV identification.

As in Leonard *et al.,* the likelihood of scenarios (i) and (ii) for a given *N_b_* are expressed as

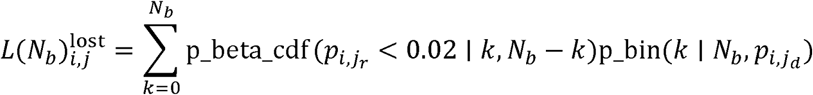

Where p_beta_cdf is the cumulative distribution function for the beta distribution. Note that if *k* = 0(i.e. the iSNV was not transmitted) then the term reduces to the probability of not drawing the variant in *N_b_* draws.

The likelihood of the variant being transmitted but not detected in the recipient given a bottleneck of *N_b_* is described by

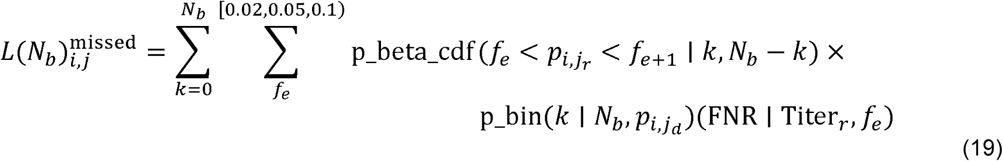

This is the likelihood of the variant existing in the ranges [0.0.2,0.05] or [0.05,0.1] given an initial frequency of *k/N_b_* and a bottleneck size of *N_b_* multiplied by the expected false negative rate (FNR) given the titer of the recipient and the lower frequency bound. As in our diffusion model, we assumed perfect sensitivity for detection of iSNV present above 10%, rounded recipient titers down to the nearest log_10_ titer (e.g. 10^3^, 10^4^, 10^5^) and assumed the entire range [*f_e_,f_e+1_*] has the same sensitivity as the lower bound.

The likelihood of *λ* for iSNV that are not observed in the recipient is then given by summing equations 18 and 19 across all possible *N_b_*.

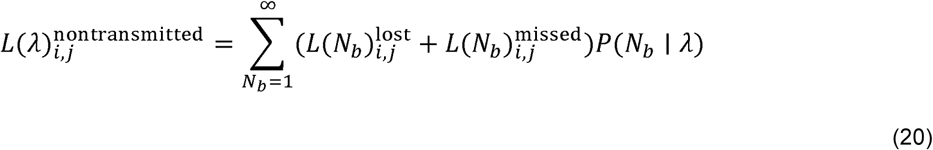

The log likelihood of the total dataset is then determined by summing log of equations 17 and 20 (as applicable) across all polymorphic sites in each donor. (As before here we sum of *N_b_* within the range [1,100].)

### Simulation

In order evaluate the fits of the two transmission models, we simulated whether or not each donor iSNV was transmitted or not. This involved converting each model to a presence absence model. In each simulation, we assigned a bottleneck from the bottleneck distribution for each transmission pair. We then determined the probability of only transmitting one allele (*A_x_* where *x* Є [1,2] as in the presence/absence model above) and the probability of transmitted both alleles (*A_1_*,*A_2_* above) for each polymorphic site.

For the presence/absence model, the probabilities for each possible outcome are given by equations 11 and 12. For the beta binomial model, the probability of only observing *A_x_* at site *i* is given by

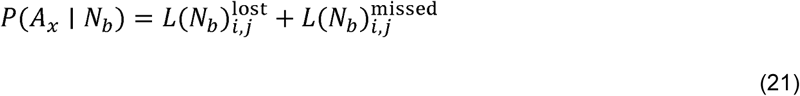

where 
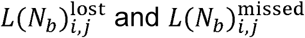
 are defined as in equations 18 and 19 respectively, but with *p_i,j_d__* replaced by 1–*p_i,j_d__*. This is simply the probability of not observing the other allele in the recipient.

Again, the probability of observing both alleles is

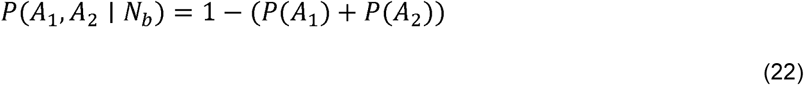

where *P*(*A_1_*)and *P*(*A_2_*)are defined as in equation 21.

### Fitting mutation rate and *N_e_*

The diffusion approximation to the Wright - Fischer model allows us to make predictions on the allele frequency spectrum of a population given a mutation rate and an effective population size. The probability of observing a mutation at frequency *p_t_* given an initial frequency of 0 can be approximated as in (Rouzine et al. 2001)

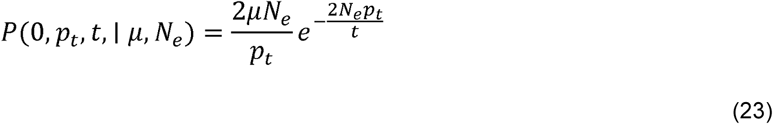

Where *μ* is the mutation rate. In this model mutation increases an allele's frequency from 0 but after that initial jump, drift is responsible for allowing the mutation to reach it's observed frequency. Because the limit of equation 23 approaches infinity as *p_t_* approaches 0 and for ease in numerical integration, we assumed that any variant present at less than 0.1% was essentially at 0%.

We then assumed each infection began as a clonal infection matching the consensus sequence observed at the time of sampling. The likelihood of observing minor alleles at the observed frequency is the given by equation 23.

As in the other within host models, we can account for nonpolymorphic sites by adding the likelihood that no mutation is present *P*(0,*p_t_ ≈ 0,t | p_t_ < 0.001,μ, N_e_*), that a mutation is present but below our level of detection *P*(0,*p_t_ ≈ 0,t | p_t_ < 0.02,μ, N_e_*), and that a mutation is present but missed due to low sensitivity at low frequencies *P*(0,*p_t_ ≈ 0,t | 0.02 < p_t_ < 0.1,μ, N_e_*). In this model we assumed 13133 mutagenic targets in each sample (the number of coding sites present in the reference strain from 2014-2015).

The probability of not observing a mutation is given by

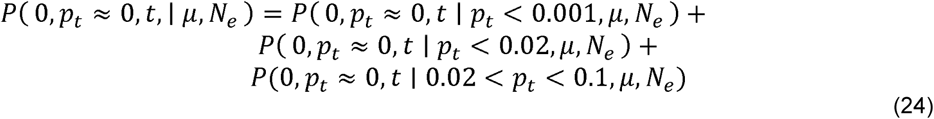

Where

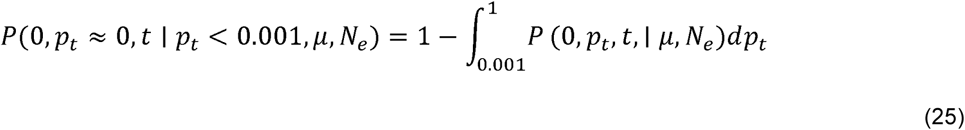

and

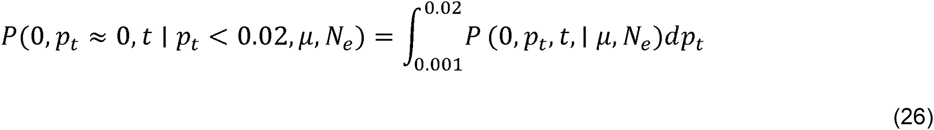

and

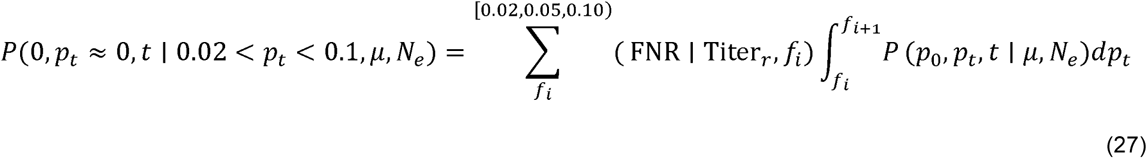

Where we follow the same convention as in equation 5 for determining the false negative rate. The log likelihood of a given *μ* and *N_e_* pair is then the sum of the log of equations 23 and 24 for all possible sites in the data set.

Annotated computer code for all analyses and for generating the figures can be accessed at https://github.com/lauringlab/Host_level_IAV_evolution

## Acknowledgments

This work was supported by a Clinician Scientist Development Award from the Doris Duke Charitable Foundation (CSDA 2013105), a University of Michigan Discovery Grant, and R01 AI118886, all to ASL. The HIVE cohort was supported by NIH R01 AI097150 and CDC U01 IP00474 to ASM. JTM was supported by the Michigan Predoctoral Training Program in Genetics (T32GM007544). RJW was supported by K08AI119182. We thank Alexey Kondrashov and Aaron King for helpful discussion.

---

**Supplementary Figure 1.** Sequence coverage for all samples. For each sample, the sliding window mean coverage was calculated using a window size of 200 and a step of 100. The distributions of these means are plotted as box plots (median, 25^th^ and 75^th^ percentiles, whiskers extend to most extreme point within median ± 1.5 x IQR) where the y-axis represents the read depth and the x-axis indicates the position of the window in a concatenated IAV genome.

**Supplementary Figure 2.** Approximate maximum likelihood trees of the concatenated coding sequences for high quality H1N1 samples. The branches are colored by season; the tip identifiers are colored by household. Arrows with numbers indicate consensus and putative minor haplotypes for samples with greater than 10 iSNV. Trees were made using FastTree.

**Supplementary Figure 3.** Approximate maximum likelihood trees of the concatenated coding sequences for high quality H3N2 samples. The branches are colored by season; the tip identifiers are colored by household. Arrows with numbers indicate consensus and putative minor haplotypes for samples with greater than 10 iSNV. Trees were made using FastTree.

**Supplementary Figure 4.** The effect of titer and vaccination on the number of iSNV identified.(A) The number of iSNV identified in an isolate (y-axis) plotted against the titer (x-axis,genomes/μl transport media). (B) The number of iSNV identified in each isolate stratified by whether that individual was vaccinated or not. Red bars indicate the median of each distribution.

**Supplementary Figure 5.** Minority nonsynonymous iSNV in global circulation. The global frequencies of the amino acids that were found as minority variants in sample isolates (x-axis) plotted overtime (y-axis). Each amino acid trace is labeled according to the H3 number scheme. All samples were isolated in December of 2014 (gray line).

**Supplementary Figure 6.** Reproducibility of iSNV identification for paired samples acquired on the same day. The x-axis represents iSNV frequencies found in the home-acquired nasal swab. The y-axis represents iSNV frequencies found the clinic-acquired combined throat and nasal swab.

**Supplementary Figure 7.** Estimate of effective bottleneck size with relaxed variant calling criteria. (A) The frequency of iSNV in both recipient and donor isolates. iSNV were identified using the original variant calling pipeline. (B) The presence-absence model fit compared to the observed data for iSNV identified using the original variant calling pipeline. The x-axis represents the frequency of donor iSNV with transmitted iSNV plotted along the top and nontransmitted iSNV plotted along the bottom. The black line indicates the probability of transmission for a given iSNV frequency as determined by logistic regression. Similar fits were calculated for 1,000 simulations with a mean bottleneck size of 2.10. Fifty percent of simulated outcomes lie in the darkly shaded region and 95% lie in the lightly shaded regions. (C) Similar to(A) but with minority iSNV identified using the current analytical framework without a frequency threshold. (D) Similar to B but with minority iSNV identified using the current analytical framework without a frequency threshold.

**Supplementary Table 1.**
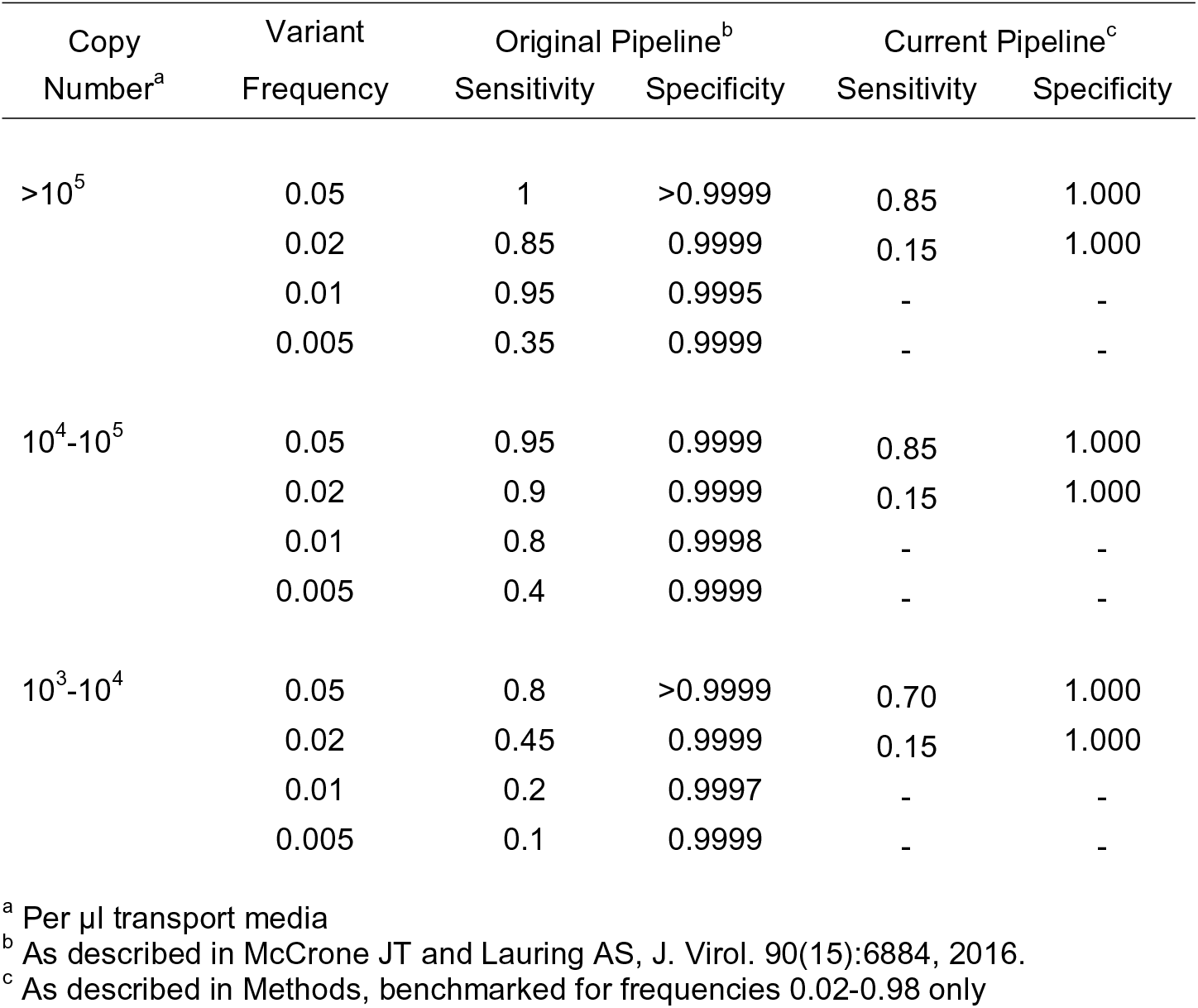
Sensitivity and specificity of variant detection.

| Copy Number<sup>a</sup> | Variant Frequency | Original Pipeline<sup>b</sup> |  | Current Pipeline<sup>c</sup> |  |
| --- | --- | --- | --- | --- | --- |
|  |  | Sensitivity | Specificity | Sensitivity | Specificity |
| >10<sup>5</sup> | 0.05 | 1 | >0.9999 | 0.85 | 1.000 |
|  | 0.02 | 0.85 | 0.9999 | 0.15 | 1.000 |
|  | 0.01 | 0.95 | 0.9995 | - | - |
|  | 0.005 | 0.35 | 0.9999 | - | - |
| 10<sup>4</sup>-10<sup>5</sup> | 0.05 | 0.95 | 0.9999 | 0.85 | 1.000 |
|  | 0.02 | 0.9 | 0.9999 | 0.15 | 1.000 |
|  | 0.01 | 0.8 | 0.9998 | - | - |
|  | 0.005 | 0.4 | 0.9999 | - | - |
| 10<sup>3</sup>-10<sup>4</sup> | 0.05 | 0.8 | >0.9999 | 0.70 | 1.000 |
|  | 0.02 | 0.45 | 0.9999 | 0.15 | 1.000 |
|  | 0.01 | 0.2 | 0.9997 | - | - |
|  | 0.005 | 0.1 | 0.9999 | - | - |

<sup>a</sup> Per µl transport media
<sup>b</sup> As described in McCrone JT and Luring AS, J. Virol. 90(15):6884, 2016.
<sup>c</sup> As described in Methods, benchmarked for frequencies 0.02-0.98 only

**Supplementary Table 2.** Nonsynonymous substitutions in HA antigenic sites.

| House ID | Enrolment ID | Symptom Onset | Subtype | Frequency | Amino Acid Change | Antigenic Site | Vaccinated | Day of Symptoms |
| --- | --- | --- | --- | --- | --- | --- | --- | --- |
| 1111 | 300481 | 3-30-2011 | H3N2 | 0.071 | E62G | E* | No | 0 |
| 2166 | 320661 | 2-13-2012 | H3N2 | 0.071 | V297A | C | Yes | 1 |
| 1302 | 301355 | 3-20-2011 | H3N2 | 0.088 | L86I | E | Yes | 1 |
| 3075 | 331045 | 12-10-2012 | H3N2 | 0.066 | I214T | D | Yes | 1 |
| 5219 | 50935 | 12-5-2014 | H3N2 | 0.175 | F193S | B*<sup>†</sup> | No | 3 |
| 5263 | 51106 | 12-6-2014 | H3N2 | 0.111 | T128A | B | Yes | 3 |
| 5290 | 51225 | 12-15-2014 | H3N2 | 0.405 | I260V | E* | Yes | 1 |
| 5302 | 51273 | 12-13-2014 | H3N2 | 0.030 | S262N | E* | Yes | 0 |
| 5098 | 50419 | 12-22-2014 | H3N2 | 0.364 | G208R | D | Yes | 4 |
| 5033 | 50141 | 12-3-2014 | H3N2 | 0.032 | A163T | B | Yes | 2 |
| 5034 | 50143 | 1-11-2015 | H3N2 | 0.119 | I307R | C | Yes | 1 |
| 5289 | 51220 | 12-13-2014 | H3N2 | 0.038 | K189N | B*<sup>†</sup> | Yes | -1 |
| 5033 | 50141 | 12-3-2014 | H3N2 | 0.025 | D53E | C* | Yes | 1 |
| 5033 | 50141 | 12-3-2014 | H3N2 | 0.023 | S312G | C | Yes | 1 |
| 5269 | 51132 | 12-6-2014 | H3N2 | 0.028 | I242T | D | Yes | 2 |
| 5147 | 50630 | 11-18-2014 | H3N2 | 0.164 | I242L | D | Yes | 1 |
| 5034 | 50143 | 1-11-2015 | H3N2 | 0.161 | I307R | C | Yes | 2 |
| 4185 | UM40738 | 12-14-2013 | H1N1 | 0.021 | R208K | Ca | No | 2 |

\* Sites observed to vary between antigenically distinct strains in Wiley et al., 1981 and Smith DJ et al., 2004.
<sup>†</sup> Sites located in the "antigenic ridge" identified in Koel et al., 2013.

**Supplementary Figure 1.**
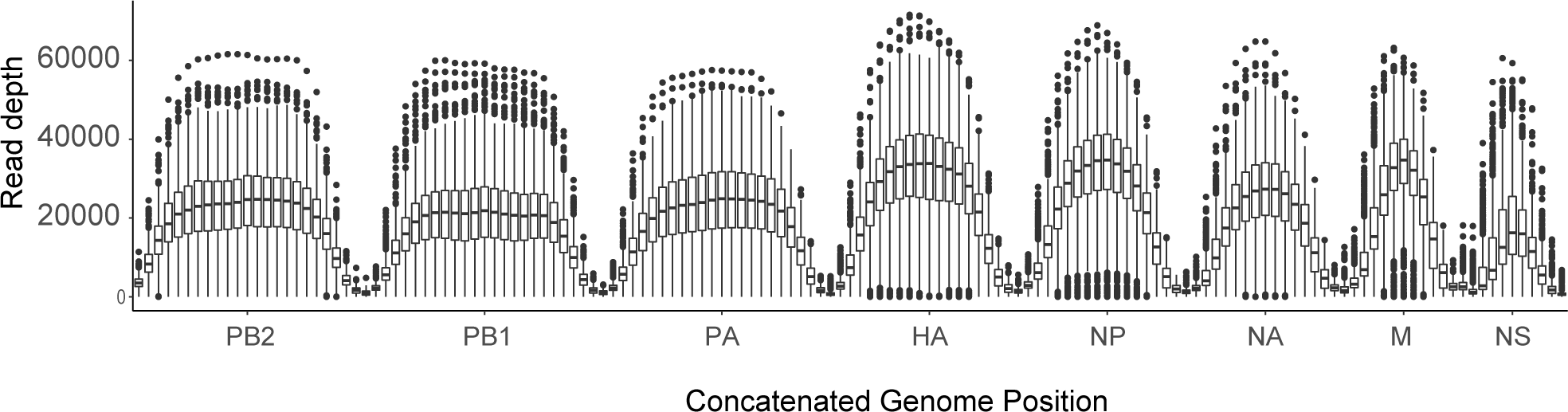
Sequence coverage for all samples. For each sample, the sliding window mean coverage was calculated using a window size of 200 and a step of 100. The distributions of these means are plotted as box plots (median, 25^th^ and 75^th^ percentiles, whiskers extend to most extreme point within median ± 1.5 x IQR) where the y-axis represents the read depth and the x-axis indicates the position of the window in a concatenated IAV genome.

**Supplementary Figure 2.**
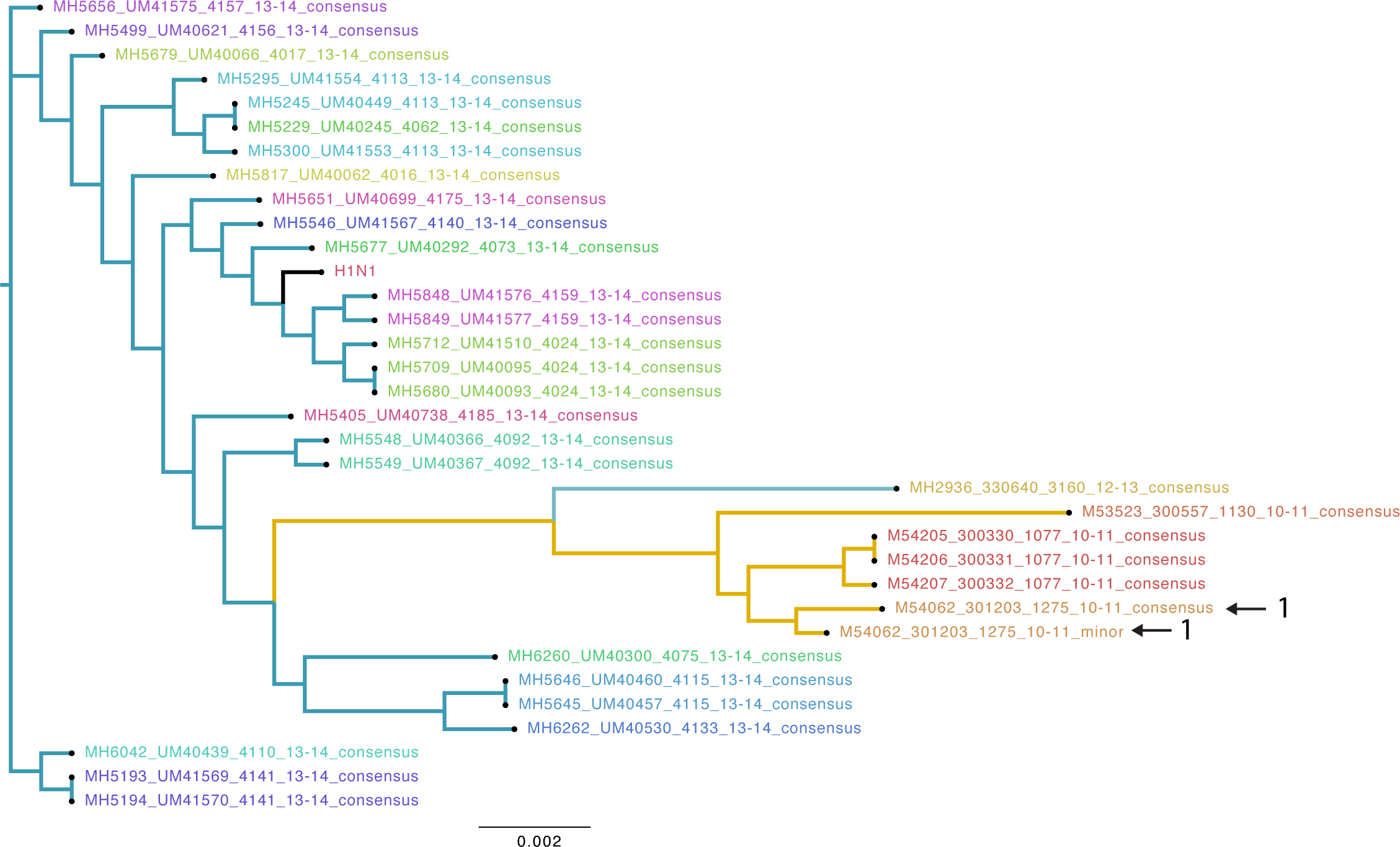
Approximate maximum likelihood trees of the concatenated coding sequences for high quality H1N1 samples. The branches are colored by season; the tip identifiers are colored by household. Arrows with numbers indicate consensus and putative minor haplotypes for samples with greater than 10 iSNV. Trees were made using FastTree.

**Supplementary Figure 3.**
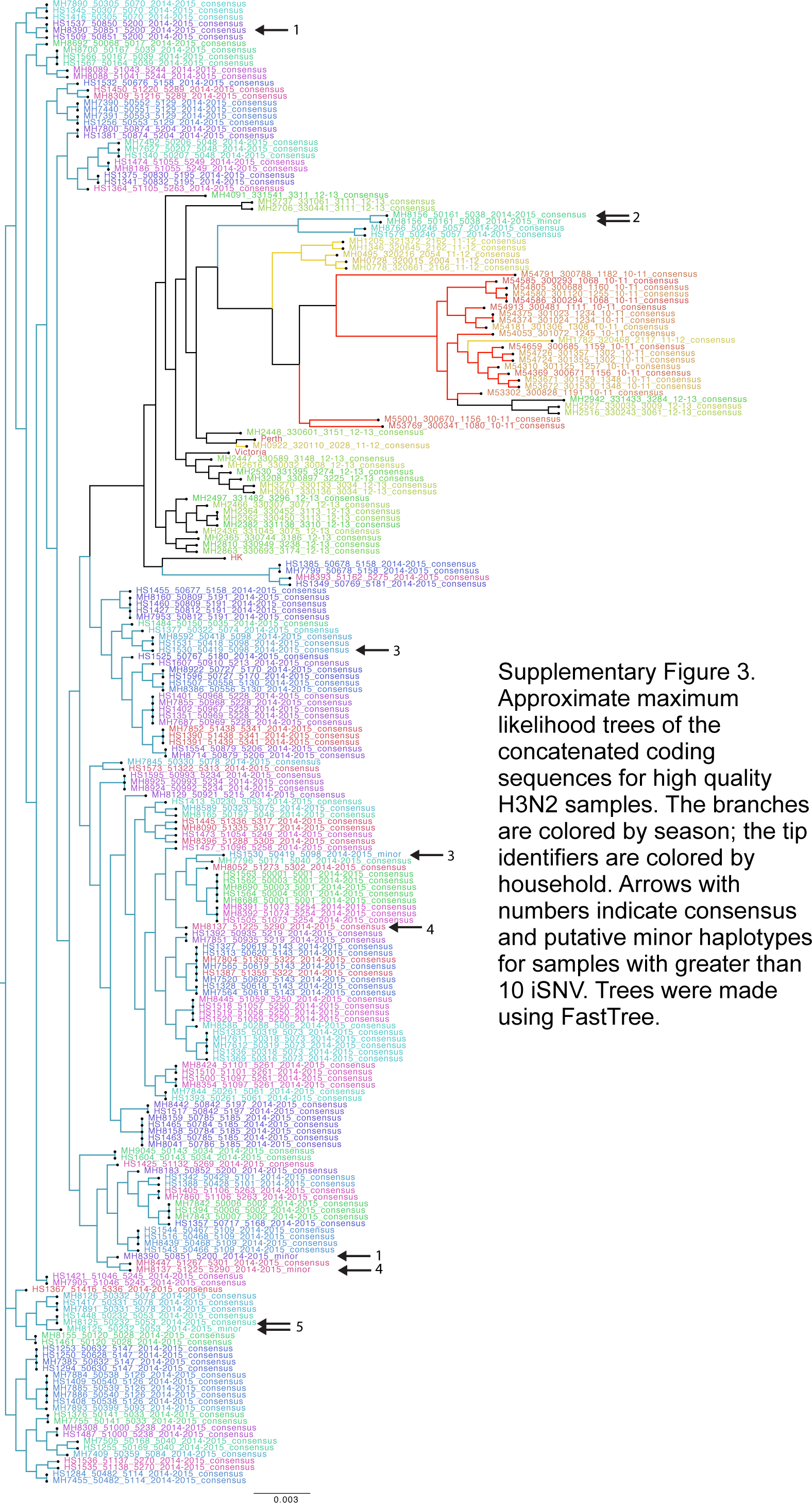
Approximate maximum likelihood trees of the concatenated coding sequences for high quality H3N2 samples. The branches are colored by season; the tip identifiers are colored by household. Arrows with numbers indicate consensus and putative minor haplotypes for samples with greater than 10 iSNV. Trees were made using FastTree.

**Supplementary Figure 4.**
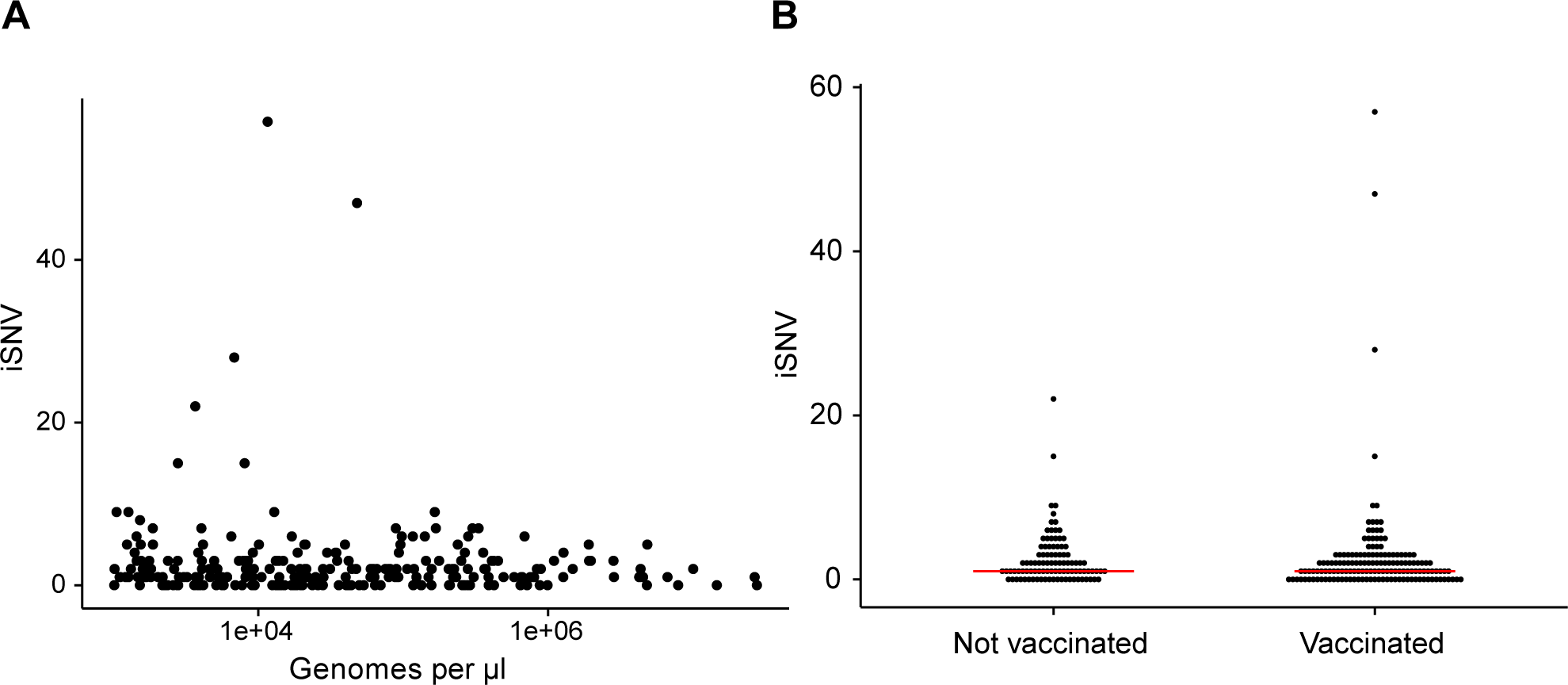
The effect of titer and vaccination on the number of iSNV identified. (A) The number of iSNV identified in an isolate (y-axis) plotted against the titer (x-axis, genomes/μl transport media). (B) The number of iSNV identified in each isolate stratified by whether that individual was vaccinated or not. Red bars indicate the median of each distribution.

**Supplementary Figure 5.**
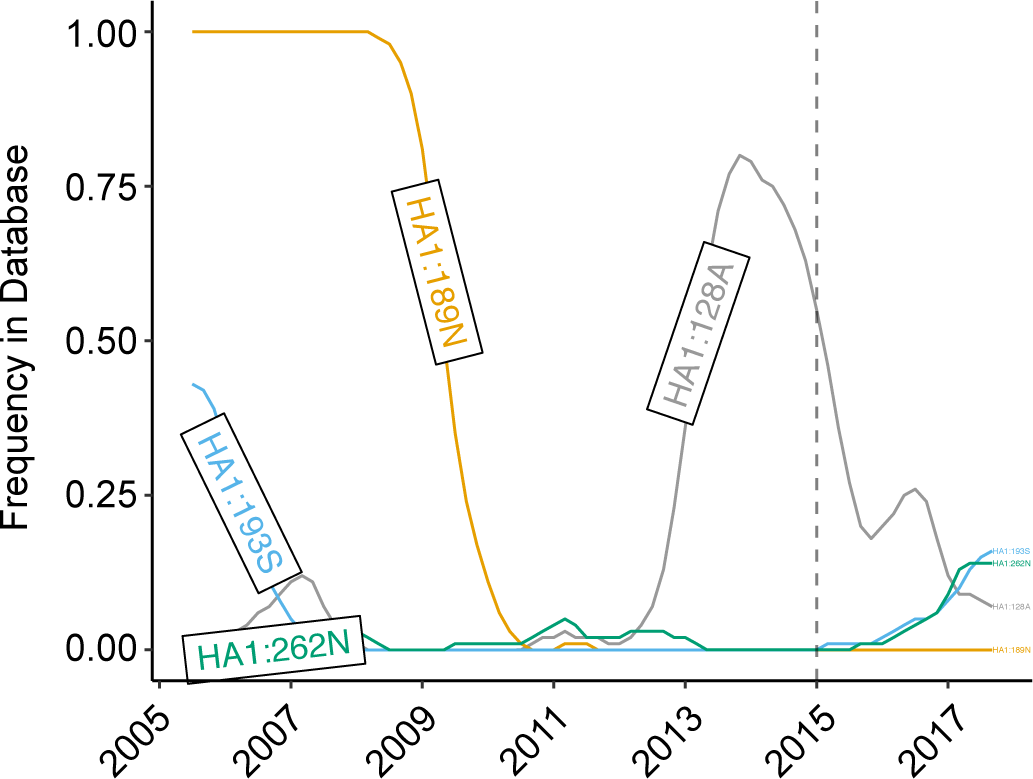
Minority nonsynonymous iSNV in global circulation. The global frequencies of the amino acids that were found as minority variants in sample isolates (x-axis) plotted overtime (y-axis). Each amino acid trace is labeled according to the H3 number scheme. All samples were isolated in December of 2014 (gray line).

**Supplementary Figure 6.**
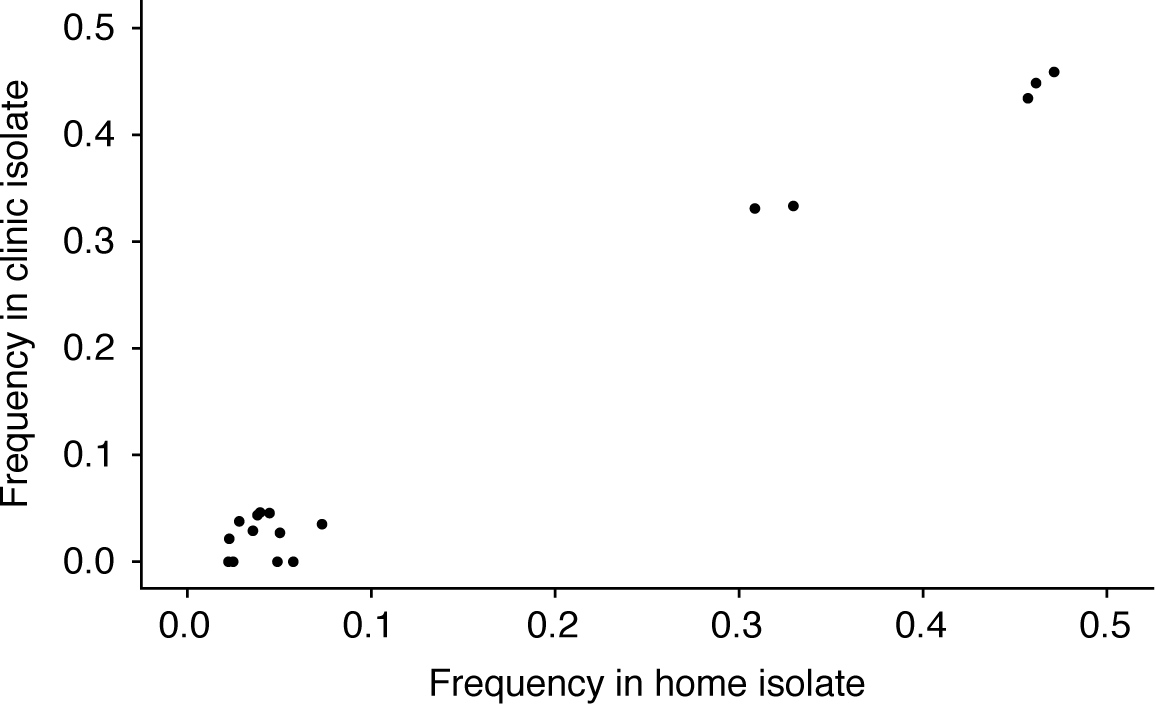
Reproducibility of iSNV identification for paired samples acquired on the same day. The x-axis represents iSNV frequencies found in the home-acquired nasal swab. The y-axis represents iSNV frequencies found the clinic-acquired combined throat and nasal swab.

**Supplementary Figure 7.**
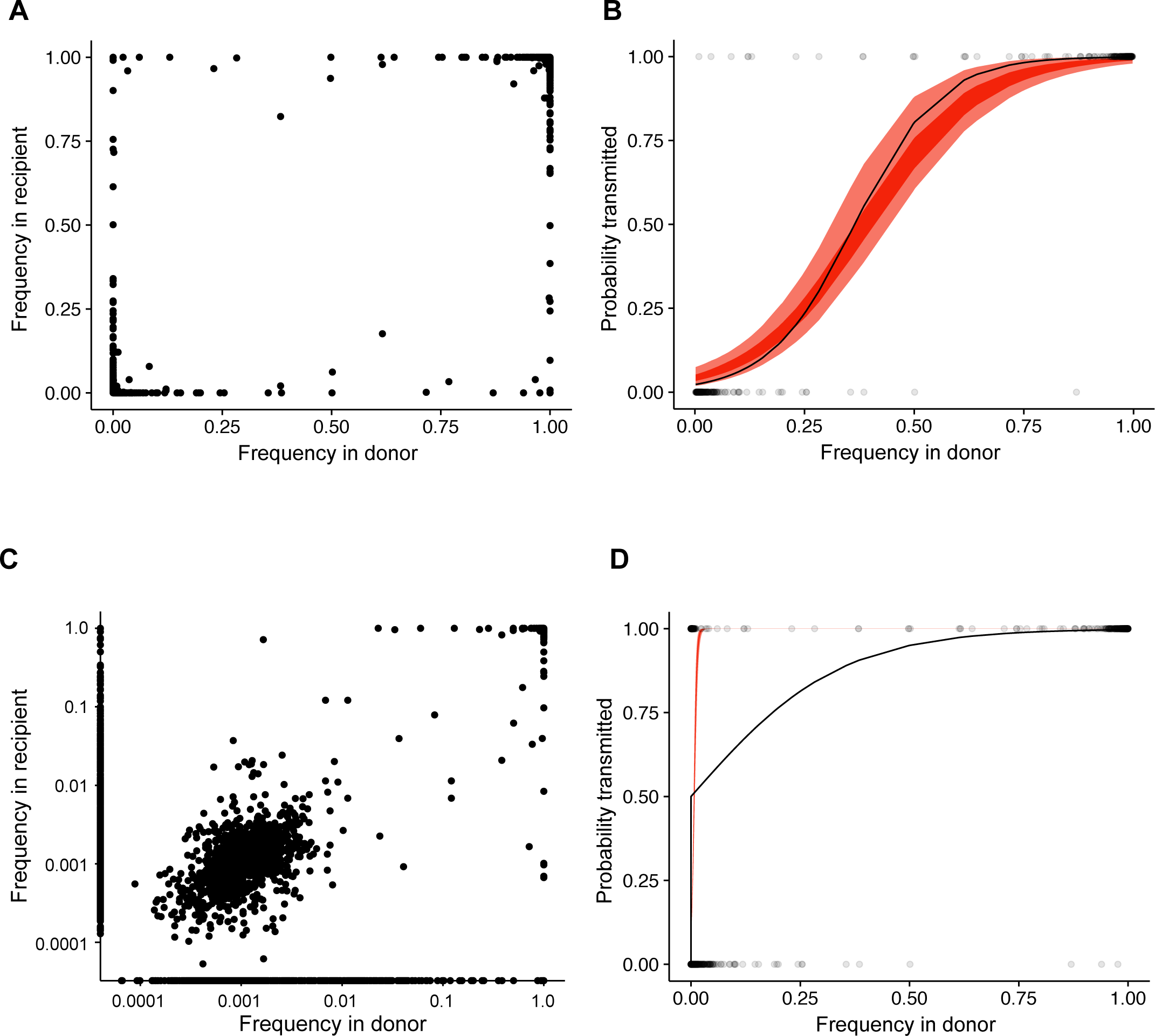
Estimate of effective bottleneck size with relaxed variant calling criteria. (A) The frequency of iSNV in both recipient and donor isolates. iSNV were identified using the original variant calling pipeline. (B) The presence-absence model fit compared to the observed data for iSNV identified using the original variant calling pipeline. The x-axis represents the frequency of donor iSNV with transmitted iSNV plotted along the top and nontransmitted iSNV plotted along the bottom. The black line indicates the probability of transmission for a given iSNV frequency as determined by logistic regression. Similar fits were calculated for 1,000 simulations with a mean bottleneck size of 2.10. Fifty percent of simulated outcomes lie in the darkly shaded region and 95% lie in the lightly shaded regions.(C) Similar to (A) but with minority iSNV identified using the current analytical framework without a frequency threshold. (D) Similar to B but with minority iSNV identified using the current analytical framework without a frequency threshold.

## References

Alizon, S., Luciani, F. & Regoes, R.R., 2011. Epidemiological and clinical consequences of within-host evolution. Trends in microbiology, 19(1), pp.24–32.

Archetti, I. & Horsfall, F.L., 1950. Persistent antigenic variation of influenza A viruses after incomplete neutralization in ovo with heterologous immune serum. The Journal of experimental medicine, 92(5), pp.441–462.

Caton, A.J. et al., 1982. The antigenic structure of the influenza virus A/PR/8/34 hemagglutinin (H1 subtype). Cell, 31(2 Pt 1), pp.417–427.

Debbink, K. et al., 2017. Vaccination has minimal impact on the intrahost diversity of H3N2 influenza viruses. PLoS pathogens, 13(1), p.e1006194.

Dinis, J.M. et al., 2016. Deep Sequencing Reveals Potential Antigenic Variants at Low Frequencies in Influenza A Virus-Infected Humans S. Schultz-Cherry, ed. Journal of Virology, 90(7), pp.3355–3365.

Doud, M.B., Hensley, S.E. & Bloom, J.D., 2017. Complete mapping of viral escape from neutralizing antibodies A. S. Lauring, ed. PLoS pathogens, 13(3), pp.e1006271–20.

Ewens, W.J., 2004. Mathematical Population Genetics 1, Springer Science & Business Media.

Flannery, B. et al., 2016. Enhanced Genetic Characterization of Influenza A(H3N2) Viruses and Vaccine Effectiveness by Genetic Group, 2014-2015. The Journal of Infectious Diseases, 214(7), pp.1010–1019.

Foll, M. et al., 2014. Influenza virus drug resistance: a time-sampled population genetics perspective. PLoS genetics, 10(2), p.e1004185.

Foll, M., Shim, H. & Jensen, J.D., 2015. WFABC: a Wright-Fisher ABC-based approach for inferring effective population sizes and selection coefficients from time-sampled data. Molecular ecology resources, 15(1), pp.87–98.

Geoghegan, J.L., Senior, A.M. & Holmes, E.C., 2016. Pathogen population bottlenecks and adaptive landscapes: overcoming the barriers to disease emergence. Proceedings. Biological sciences / The Royal Society, 283(1837), pp.20160727–9.

Gerstung, M. et al., 2012. Reliable detection of subclonal single-nucleotide variants in tumour cell populations. Nature Communications, 3, p.811.

Ghedin, E. et al., 2011. Deep sequencing reveals mixed infection with 2009 pandemic influenza A (H1N1) virus strains and the emergence of oseltamivir resistance. The Journal of Infectious Diseases, 203(2), pp.168–174.

Gubareva, L.V. et al., 2001. Selection of influenza virus mutants in experimentally infected volunteers treated with oseltamivir. The Journal of Infectious Diseases, 183(4), pp.523–531.

Herfst, S. et al., 2012. Airborne transmission of influenza A/H5N1 virus between ferrets. Science (New York, NY), 336(6088), pp.1534–1541.

Holmes, E.C., 2009. RNA virus genomics: a world of possibilities. The Journal of clinical investigation, 119(9), pp.2488–2495.

Hughes, J. et al., 2012. Transmission of equine influenza virus during an outbreak is characterized by frequent mixed infections and loose transmission bottlenecks. PLoS pathogens, 8(12), p.e1003081.

Iqbal, M. et al., 2009. Within-host variation of avian influenza viruses. Philosophical transactions of the Royal Society of London Series B, Biological sciences, 364(1530), pp.2739–2747.

Kao, R.R. et al., 2014. Supersize me: how whole-genomesequencing and big data aretransforming epidemiology. Trends in microbiology, pp.1–10.

Kimura, M., 1955. SOLUTION OF A PROCESS OF RANDOM GENETIC DRIFT WITH A CONTINUOUS MODEL. Proceedings of the National Academy of Sciences of the United States of America, 41(3), pp.144–150.

Kimura, M., 1971. Theoretical foundation of population genetics at the molecular level. Theoretical population biology, 2(2), pp.174–208.

Kimura, M. & Ohta, T., 1969. The Average Number of Generations until Fixation of a Mutant Gene in a Finite Population. Genetics, 61(3), pp.763–771.

Koel, B.F. et al., 2013. Substitutions near the receptor binding site determine major antigenic change during influenza virus evolution. Science (New York, NY), 342(6161), pp.976–979.

Kouyos, R.D., Althaus, C.L. & Bonhoeffer, S., 2006. Stochastic or deterministic: what is the effective population size of HIV-1? Trends in microbiology, 14(12), pp.507–511.

Kugelman, J.R. et al., 2017. Error baseline rates of five sample preparation methods used to characterize RNA virus populations K. K. Tee, ed., 12(2), pp.e0171333–13. Available at: http://dx.plos.org/10.1371/journal.pone.0171333.

Lee, M.-S. & Chen, J.S.-E., 2004. Predicting antigenic variants of influenza A/H3N2 viruses. Emerging infectious diseases, 10(8), pp.1385–1390.

McCrone, J.T. & Lauring, A.S., 2016. Measurements of Intrahost Viral Diversity Are Extremely Sensitive to Systematic Errors in Variant Calling. Journal of Virology, 90(15), pp.6884–6895.

Monto, A.S. et al., 2014. Frequency of acute respiratory illnesses and circulation of respiratory viruses in households with children over 3 surveillance seasons. The Journal of Infectious Diseases, 210(11), pp.1792–1799.

Murcia, P.R. et al., 2010. Intra‐ and interhost evolutionary dynamics of equine influenza virus. Journal of Virology, 84(14), pp.6943–6954.

Myers, S., Fefferman, C. & Patterson, N., 2008. Can one learn history from the allelic spectrum? Theoretical population biology, 73(3), pp.342–348.

Neher, R.A. & Bedford, T., 2015. nextflu: real-time tracking of seasonal influenza virus evolution in humans. Bioinformatics (Oxford, England), p.btv381.

Nelson, M.I. & Holmes, E.C., 2007. The evolution of epidemic influenza. Nature Reviews Genetics, 8(3), pp.196–205.

Nelson, M.I. et al., 2006. Stochastic processes are key determinants of short-term evolution in influenza a virus. PLoS pathogens, 2(12), p.e125.

Ohmit, S.E. et al., 2015. Influenza vaccine effectiveness in households with children during the 2012-2013 season: assessments of prior vaccination and serologic susceptibility. The Journal of Infectious Diseases, 211(10), pp.1519–1528.

Ohmit, S.E. et al., 2013. Influenza Vaccine Effectiveness in the Community and the Household. Clinical infectious diseases: an official publication of the Infectious Diseases Society of America, 56(10), pp.1363–1369.

Ohmit, S.E. et al., 2016. Substantial Influenza Vaccine Effectiveness in Households With Children During the 2013-2014 Influenza Season, When 2009 Pandemic Influenza A(H1N1) Virus Predominated. The Journal of Infectious Diseases, 213(8), pp.1229–1236.

Pauly, M.D., Procario, M.C. & Lauring, A.S., 2017. A novel twelve class fluctuation test reveals higher than expected mutation rates for influenza A viruses. eLife, 6, p.e26437.

Peck, K.M., Chan, C.H.S. & Tanaka, M.M., 2015. Connecting within-host dynamics to the rate of viral molecular evolution. Virus Evolution, 1(1), p.vev013.

Petrie, J.G. et al., 2017. Application of an Individual-Based Transmission Hazard Model for Estimation of Influenza Vaccine Effectiveness in a Household Cohort. American journal of epidemiology.

Petrie, J.G. et al., 2013. Influenza Transmission in a Cohort of Households with Children: 2010-2011 J. McVernon, ed. PLoS ONE, 8(9), p.e75339.

Petrova, V.N. & Russell, C.A., 2017. The evolution of seasonal influenza viruses. Nature Reviews Microbiology, 2, p.517.

Poon, L.L.M. et al., 2016. Quantifying influenza virus diversity and transmission in humans. Nature Genetics, 48(2), pp.195–200.

Rambaut, A. et al., 2008. The genomic and epidemiological dynamics of human influenza A virus. Nature, 453(7195), pp.615–619.

Rogers, M.B. et al., 2015. Intrahost dynamics of antiviral resistance in influenza A virus reflect complex patterns of segment linkage, reassortment, and natural selection. mBio, 6(2).

Rouzine, I.M., Rodrigo, A. & Coffin, J.M., 2001. Transition between stochastic evolution and deterministic evolution in the presence of selection: general theory and application to virology. Microbiology and molecular biology reviews: MMBR, 65(1), pp.151–185.

Russell, C.A. et al., 2012. The potential for respiratory droplet-transmissible A/H5N1 influenza virus to evolve in a mammalian host. Science (New York, NY), 336(6088), pp.1541–1547.

Sanjuán, R. et al., 2010. Viral mutation rates. Journal of Virology, 84(19), pp.9733–9748.

Smith, D.J. et al., 2004. Mapping the antigenic and genetic evolution of influenza virus. Science (New York, NY), 305(5682), pp.371–376.

Sobel Leonard, A. et al., 2016. Deep Sequencing of Influenza A Virus from a Human Challenge Study Reveals a Selective Bottleneck and Only Limited Intrahost Genetic Diversification D. S. Lyles, ed. Journal of Virology, 90(24), pp.11247–11258.

Sobel Leonard, A. et al., 2017. Transmission Bottleneck Size Estimation from Pathogen Deep‐ Sequencing Data, with an Application to Human Influenza A Virus. Journal of Virology, 91(14).

Varble, A. et al., 2014. Influenza A virus transmission bottlenecks are defined by infection route and recipient host. Cell Host and Microbe, 16(5), pp.691–700.

Visher, E. et al., 2016. The Mutational Robustness of Influenza A Virus N. M. Ferguson, ed. PLoS pathogens, 12(8), pp.e1005856–25.

Wiley, D.C., Wilson, I.A. & Skehel, J.J., 1981. Structural identification of the antibody-binding sites of Hong Kong influenza haemagglutinin and their involvement in antigenic variation. Nature, 289(5796), pp.373–378.

Wilker, P.R. et al., 2013. Selection on haemagglutinin imposes a bottleneck during mammalian transmission of reassortant H5N1 influenza viruses. Nature Communications, 4, p.2636.

Williamson, E.G. & Slatkin, M., 1999. Using maximum likelihood to estimate population size from temporal changes in allele frequencies. Genetics, 152(2), pp.755–761.

Xu, R. et al., 2010. Structural basis of preexisting immunity to the 2009 H1N1 pandemic influenza virus. Science (New York, NY), 328(5976), pp.357–360.

Xue, K.S. et al., 2017. Parallel evolution of influenza across multiple spatiotemporal scales. eLife, 6, p.e26875.

Zanini, F. et al., 2017. In vivo mutation rates and the landscape of fitness costs of HIV-1. Virus Evolution, 3(1), p.vex003.

Zhou, B. et al., 2009. Single-reaction genomic amplification accelerates sequencing and vaccine production for classical and Swine origin human influenza a viruses. Journal of Virology, 83(19), pp.10309–10313.

Zwart, M.P. & Elena, S.F., 2015. Matters of Size: Genetic Bottlenecks in Virus Infection and Their Potential Impact on Evolution. Annual Review of Virology, 2(1), pp.161–179.

